# Spatial analysis of the metastatic brain tumor immune and extracellular matrix microenvironment

**DOI:** 10.1101/2022.08.30.505945

**Authors:** Samuel S. Widodo, Marija Dinevska, Lucero Cuzcano, Michael Papanicolaou, Thomas R. Cox, Stanley S. Stylli, Theo Mantamadiotis

**Author notes:** Equal contribution (Samuel S. Widodo, Marija Dinevska).

## Abstract

Metastatic cancer is responsible for the overwhelming majority of cancer-related deaths with metastatic tumors being the most common neoplasms affecting the central nervous system. One of the major factors regulating tumor biology is the tumor microenvironment. However, little is known about the cellular and non-cellular composition of metastatic brain tumors and how tumor cell ontogeny influences the metastatic brain tumor microenvironment. By integrating multiplex immunohistochemistry and histopathological analysis to investigate composition and the spatial relationship between neoplastic cells, infiltrating and brain resident immune cells and the extracellular matrix, we demonstrate that metastatic brain tumors exhibit differences in ECM deposition, compared with the most common primary brain tumor type, glioblastoma, and that the dominant immune cell types in metastatic brain tumors are immunosuppressive macrophages, which preferentially localize to ECM-rich stromal regions.

## Introduction

Intracranial metastases are the most frequent type of brain tumor and one of the most common neurological complications of systemic cancer, representing about 20% of intracranial tumors diagnosed in adults [1]. An estimated 40% of all patients with cancer will develop brain metastases (BrM), which are a common feature of advanced disease. The two most common types of primary tumors metastasizing to the brain are lung cancer, accounting for almost half of all BrM, followed by breast cancer accounting for 15-30% of metastatic brain cancer [2]. At the time of diagnosis, the majority of BrM patients harbor multiple lesions, and most will experience some degree of neurocognitive impairment during the course of the disease. The median survival for untreated patients is 1 to 2 months, and about 6 months for patients who have undergone various combinations of surgery, chemotherapy, and radiotherapy [3]. As a direct cause of morbidity and mortality in cancer patients, BrM remains one of the most challenging diseases in clinical oncology. Other than radiotherapy, to date there is no standard treatment used for patients with BrM. The discovery of mechanisms regulating BrM biology, leading to the implementation of new therapeutic approaches for the treatment of patients with BrM to improve patient outcome is of paramount importance.

One of the major factors regulating BrM biology is the tumor microenvironment (TME), which is composed of both cellular and non-cellular factors. Despite the promising outcomes of clinical trials investigating the efficacy of immunotherapies in several metastatic cancers, there is limited data available on the impact of these therapies on the BrM tumor microenvironment. This is particularly pertinent for the central nervous system, which has a unique immunological microenvironment compared to other metastatic tumor host organs [4]. Past studies have shown that the number of infiltrating immune cells in BrM differs from immune cell infiltration in the major primary brain cancer, glioblastoma (GBM), and varies according to primary tumor ontogeny. It has been shown that melanoma BrM tumors are characterized by a higher number of CD8+ T-cells compared to GBM [5]. BrMs also show lower microglial infiltration compared to GBM, but exhibit substantial infiltration of other myeloid cells, including protumorigenic immunosuppressive bone marrow-derived macrophages [6].

Moreover, BrM tumor cells interact with brain resident cells, other than microglia, including neurons and astrocytes, which play roles in supporting BrM growth and colonization, by secreting pro-tumorigenic cytokines and chemokines, which also contribute to chemotherapy resistance in BrM [7] [8].

Another relatively unexplored property of the BrM TME, is the deposition and organization of the extracellular matrix (ECM), which is the major component of the matrisome, and includes a multitude of factors, which include a spectrum of ECM modifying enzymes, known as the matrix metalloproteinases (MMPs). A feature of many primary cancers is the desmoplastic reaction which is accompanied by an increase in collagen deposition and tissue remodeling, especially in tissues in which collagen is a substantial component of the healthy tissue. However, the healthy human brain parenchyma exhibits only low levels of ECM proteins, including the major ECM protein type, collagen [9]. Further, BrM cells can facilitate the compositional makeup of the brain tumor ECM, by influencing mRNA expression of growth factors, MMPs and ECM proteins directly, including collagen and fibronectin [10] [11].

Despite the appreciation that ECM proteins regulate tumorigenesis, the role of the ECM in promoting BrM biology remains unclear. Thus, much of the current understanding of the role of the ECM in regulating the growth of BrM is drawn from existing knowledge of the primary tumors. For example, there is a positive correlation between ECM stiffness, tumor invasion and resistance to therapy in breast cancer [12]. Moreover, in vitro experiments show that collagen, a major ECM component in many tumor types, induces immunosuppressive macrophage polarization [13]. This has also been shown in breast cancer, where ECM stiffening correlates with tumor infiltration of pro-tumorigenic CD68^+^ CD163^+^ macrophages [12]. Expression of additional ECM components, including hyaluronan and collagen are also associated with poor prognosis in primary cancers [14] [15] [16] [17].

Despite the early encouraging outcomes observed with the use of immunotherapeutic based approaches for the treatment of non-central nervous system cancers, in addition to the limited clinical trials conducted for BrM patients with lung cancer and melanoma metastases [18], the response is transient and increases in median survival for these patients remains modest. To date, investigations into metastatic cancer biology have focused on the molecular and cellular characteristics of the metastatic cells and the mechanisms regulating their metastatic transition. Several genomic studies have shed light on the molecular and genetic properties driving metastasis of primary tumor cells to specific host organs, including the brain [19–21]. For example, in a breast cancer mouse model, analysis of the ECM in several metastatic host tissue niches of breast cancer xenografts, demonstrated that the brain is characterized by fewer but more diverse and unique matrisomal proteins, compared to other metastatic host tissues [22]. Detailed properties of organ-specific matrisomes in metastatic niches, including the brain, are extensively reviewed by Deasy and Erez [23].

By integrating multispectral immunohistochemistry (mIHC) and histopathological analysis of BrM tissue to interrogate the spatial relationship between the ECM and cellular tumor components, we examined the ECM content, focusing on collagen deposition, and the corresponding localization of T-cells and myeloid cells. We show that metastatic brain tumors exhibit substantial ECM deposition, which differs from primary brain tumors, and that immune cells are enriched in ECM-stromal regions. We also show that the majority of bone marrow derived macrophages are immunosuppressive.

## Methods

### Tumor tissue

All metastatic brain tumor tissue used in this study was localized within the cerebrum or cerebellum. Details of patient samples are shown in Supplementary Table 1. All brain metastatic and GBM tissue specimens were formalin-fixed paraffin embedded (FFPE) and sourced from US Biomax (GL806f and GL861a). All metastatic brain tumor tissue was collected using HIPPA (Health Insurance Portability and Accountability Act USA) approved protocols and donor informed consent. For GBM tissue, human ethics approval was covered by project application 1853511 and was approved by the Medicine and Dentistry Human Ethics Sub-Committee, The University of Melbourne.

**Table 1.** Macrophage subset biomarker identification

| Macrophage subset | Phenotype |
| --- | --- |
| CD68<sup>+</sup> HLA-DR<sup>+</sup> | Pro-inflammatory macrophage |
| CD68<sup>+</sup> CD206<sup>+</sup> | Anti-inflammatory macrophage |
| CD68<sup>+</sup> CD163<sup>+</sup> |  |
| CD68<sup>+</sup> CD163<sup>+</sup> CD206<sup>+</sup> |  |
| CD68<sup>+</sup> HLA-DR<sup>+</sup> (CD163<sup>+</sup> and/or CD206<sup>+</sup>) | Mixed phenotype |
| CD68<sup>+</sup> | Undefined |

### Automated multiplex immunohistochemistry staining

Multiplex IHC staining was automated using the Bond RX stainer (Leica Biosystems) with the Opal 7-Color IHC kit (Akoya Biosciences). 4μm thick FFPE tissue sections were baked at 60°C for 1 hour prior to deparaffinization in xylene and rehydration in a series of graded ethanol. Heat-induced antigen retrieval was performed using either a Citrate pH 6 buffer or Tris-Ethylenediaminietetraacetic acid (EDTA) pH 9 buffer. Endogenous peroxidase activity was blocked with 3% hydrogen peroxide prior to the addition of any primary antibodies which were prepared in the Opal Blocking/Antibody Diluent (Akoya biosciences). A pan-immune antibody panel and a macrophage subset antibody panel were used to investigate immune cell localization in brain metastasis samples. The pan-immune panel included CD68 (Abcam ab955, 1:100), TMEM119 (Abcam 185333, 1:1000), CD11c (Abcam ab52632, 1:1000) and CD3 (Abcam ab16669, 1:150). The macrophage subset antibody panel included CD68 (Abcam ab955, 1:100), CD163 (Cell Signaling Technology #93498, 1:250), CD206 (Abcam ab64693, 1:2000), HLA-DR (Cell Signaling Technology #97971, 1:100), Cytokeratin (AE1/AE3) (Dako M3515, 1:400) and TMEM119 (Abcam 185333, 1:1000). All tissue slides were incubated with a primary antibody for 1 hour at room temperature prior to the incubation with the Opal Polymer HRP Ms + Rb secondary antibody for 30 minutes. Immunofluorescence was visualized using Opal fluorophores (Opal 520, 540, 570, 620, 650 and 690) diluted at 1:150 in a Plus Automation Amplification Diluent (Akoya biosciences). Serial multiplexing was performed by repeating the sequence of antigen retrieval, primary antibody and Opal polymer incubation, followed by Opal fluorophore visualization for all six antibodies. Tissue sections were counterstained with DAPI, and cover-slipped using ProLong Glass Antifade Mountant.

### Multi-spectral image acquisition and image analysis

Multispectral images were acquired using the Vectra 3.0.5 Multispectral Imaging Platform (Perkin Elmer, USA) at 40x magnification. Spectral deconvolution was then performed using inForm 2.4.8 software (Perkin Elmer, USA) and multispectral images were fused using Halo software (Indica Labs, USA). Image analysis (cell phenotyping) was performed using the analysis algorithm, Highplex FL, where the nuclear detection sensitivity and the minimum intensity threshold for positive cell detection based on staining localization (nuclear, cytoplasmic or membrane) were defined. Spatial analysis was performed with Halo by annotating ECM-rich tissue regions positive for Masson’s Trichrome staining. CD3^+^, CD11C^+^, TMEM119^+^ and CD68^+^ macrophage subsets were counted in ECM and neoplastic cell-rich regions. Cell localization was also identified by measuring the infiltration distances of various cell types from ECM-rich regions into neoplastic cell-rich regions using the infiltration analysis module in Halo.

### Masson’s Trichrome staining

Masson’s Trichrome staining was performed as previously described [24]. Tissue slides were de-cover slipped and incubated with Carazzi hematoxylin, followed by incubation with 1% Briebrich scarlet −1% acid fuchsin solution. Tissue sections were then decolorized with 1% phosphotungstic acid and incubated with 1% Light Green, prior to a brief rinse with two changes of ethanol and xylene. The Masson’s Trichrome stained slides were then scanned using a Vectra 3.0.5 Multispectral Imaging Platform (Perkin Elmer, USA), at 10x magnification. To determine the amount of collagen (green) in each tissue core, the Tissue Classifier add-on in Halo was used. The add-on uses a machine learning algorithm to identify tissue types based on user input training and for this study, collagen-rich regions were used to train the software.

### Picrosirius Red staining

Picrosirius red staining was performed, as previously described [25]. Tumor tissue slides were stained with Picrosirius red solution for 60 minutes. Slides were then washed with an acetic acid solution, followed by rinsing in absolute ethanol before being dehydrated and cover slipped.

Images were captured using an Olympus BX53 microscope, at 4x magnification. Quantitative image analysis was performed using Fiji ImageJ software with a custom script, available at *https://github.com/TCox-Lab*, to identify and capture red, green and yellow birefringence signals [26].

### Statistical analysis

Data was analyzed using a non-parametric Kruskal-Wallis test (α=0.05), followed by a Dunn’s test for multiple group comparison, a two-tailed unpaired Student’s t-test and Wilcoxon matched-pairs signed rank test, an ordinary one-way ANOVA and Mann-Whitney test with GraphPad Prism v.9 software, as indicated in each figure legend. Correlation analysis was performed using a non-parametric Spearman’s rank correlation analysis. Statistical significance is represented by *(p<0.05), **(p<0.01), ***(p<0.001) and ****(p<0.0001). Box plots indicate median and interquartile range (IQR), as indicated in the figure legends. All statistical analysis was performed using GraphPad Prism version 9.0 (GraphPad Software, La Jolla California USA, www.graphpad.com).

## Results

### Collagen deposition and density in BrM tumors

To determine the extent of collagen deposition in BrM, tumor tissues were stained with Masson’s Trichrome stain (Figure 1A). GBM tissue was also examined to compare differences in collagen deposition between primary brain tumors and BrMs originating from lung and breast primary tumors. Analysis of collagen content indicated that collagen was significantly more abundant in GBM and BrM tumor cores compared to non-tumour tissues (Figure 1B). BrM tumor tissue cores exhibited higher collagen content, compared to GBM. There was no statistically significant difference in collagen deposition between BrM tissues across the different primary tumors, including lung, breast, skin, and thyroid (Figure 1C). To characterize the nature of the collagen, BrM tissue, GBM tissue and non-tumor tissue, were stained with Picrosirius Red, which stains all collagens, and when stained tissue with collagen is viewed using contrast-enhancing polarized light microscopy, only fibrillar collagens cause birefringence and are visible [27]. Differences in birefringence were detected as colors, ranging between red and green light, differentiating collagen fiber density and orientation. Collagen in BrM exhibited significantly higher red birefringence, indicative of thicker, denser collagen bundles, compared to GBM, while GBM tumor tissue showed a corresponding increase in yellow and green birefringence, indicative of thinner, less dense fibrillar collagens (Figure 1D-E). No significant differences in red, yellow or green birefringence were observed between lung and breast BrM ECM (Figure S1A-B).

**Figure 1.**
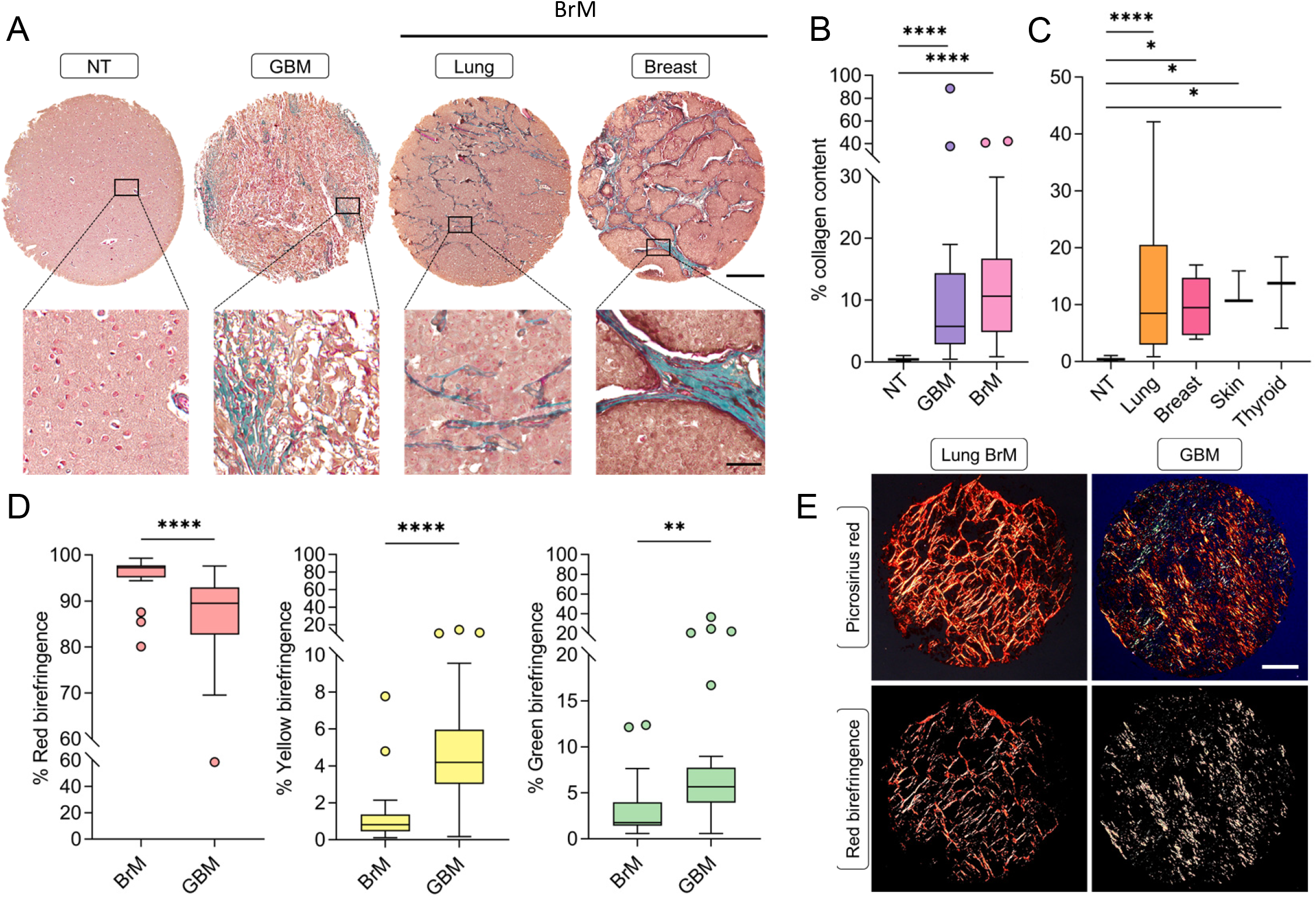
Collagen density and deposition in brain metastasis. (**A**) Representative images of Masson’s Trichrome staining showing collagen in lung, breast, skin and thyroid BrM, GBM and non-tumor tissue. Scale bars, 300μm. (**B**) Percentage of collagen deposition in non-tumor (n=15), GBM (n=35) and BrM (n=28) tissues. Collagen content was identified using the Tissue Classifier Add-on in Halo (see Methods). (**C**) Percentage of collagen deposition in non-tumor tissue (n=15), lung (n=17), breast (n=5), skin (n=3) and thyroid (n=3) BrM. (**D**) Percentage of red, yellow and green birefringence signal in BrM (n=17) and GBM (n=30) tissues. (**E**) Representative polarized light images and red birefringence signal of BrM and GBM tumor tissue. Scale bar, 300μm. All results are represented as median +/− IQR. Statistical significance was determined by Kruskal-Wallis test, followed by Dunn’s test for multiple comparisons (α=0.05) for comparison of the percentage of collagen deposition in tumor samples. Statistical significance of the birefringence signal (red, yellow and green) was determined by Mann-Whitney test. Significance is represented by *(p<0.05), **(p<0.01), ***(p<0.001) and ****(p<0.0001).

### Bone marrow-derived macrophages are the major infiltrating immune cell type in BrM

To investigate the extent of immune cell infiltration in BrM, mIHC was performed using antibodies identifying specific immune cell type biomarkers: CD68 (macrophages), CD3 (T-cells), TMEM119 (microglia), and CD11c (dendritic cells) (Figure 2A).

**Figure 2.**
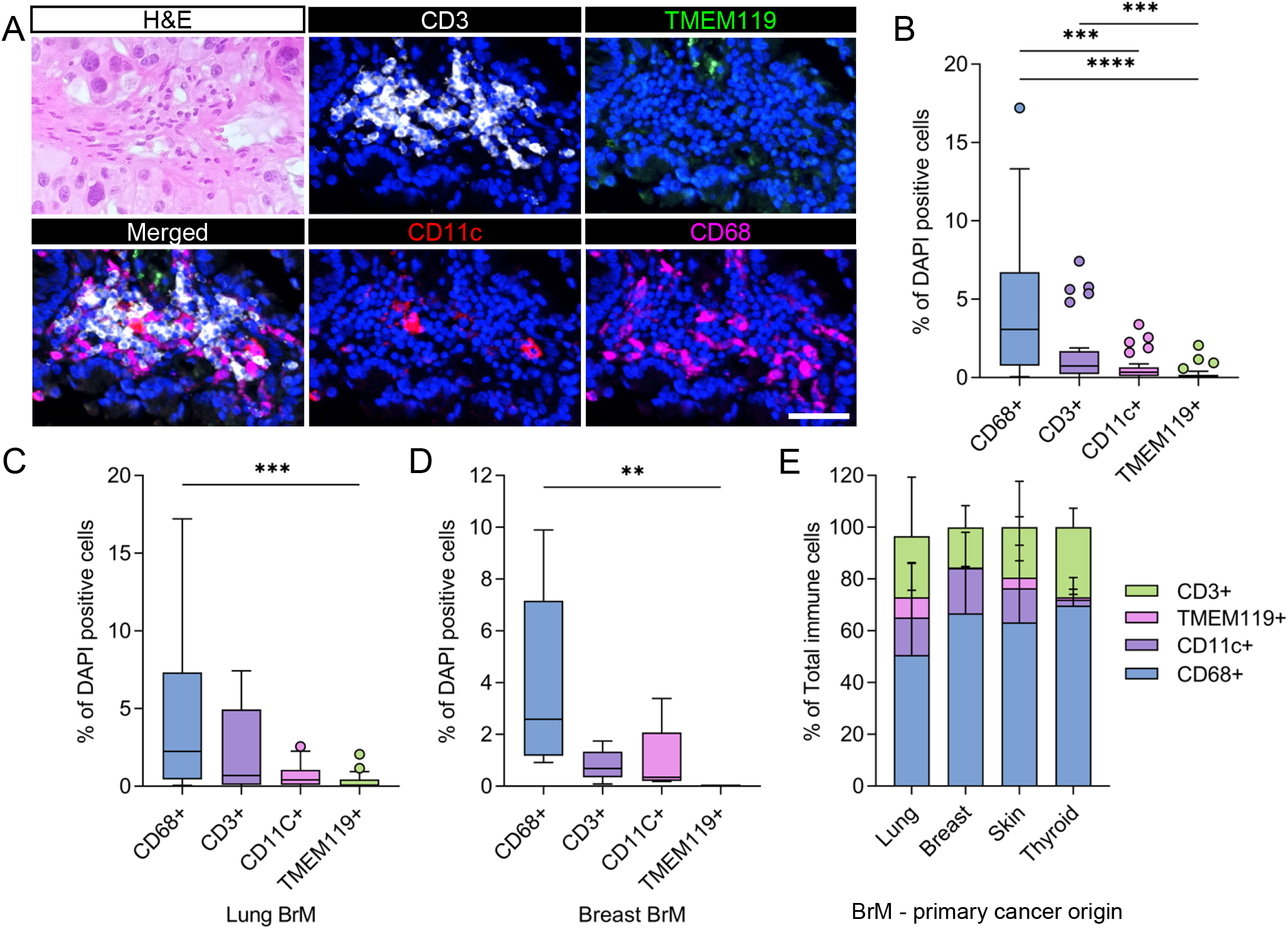
Macrophages are the major infiltrating immune cell type in BrM tumors. (**A**) Representative mIHC stained images labeled with CD68, CD11c, TMEM119 and CD3 antibodies and H&E-stained lung BrM tumor tissue. Scale bar, 50μm. (**B**) CD68^+^, CD11c^+^, TMEM119^+^ and CD3^+^ immune cell proportion relative to the total number of DAPI^+^ cells per tissue was measured. The data was derived by analyzing n=29 BrM tissue samples. CD68^+^, CD11c^+^, TMEM119^+^ and CD3^+^ immune cell proportion relative to the total number of DAPI^+^ cells per tissue was measured across BrM from (**C**) lung (n=18) and (**D**) breast primary (n=5) tumors. All results are presented as median +/− IQR. Statistical significance was determined by Kruskal-Wallis test, followed by Dunn’s test for multiple comparisons (α=0.05) for comparison between immune cell types. (**E**) Proportion of CD68^+^, CD11c^+^, TMEM119^+^ and CD3^+^ immune cells from the total number of immune cells per BrM tissue in lung (n=18), breast (n=5), skin (n=3) and thyroid (n=3) primary tumors. Results are presented as mean +/− SD. All significance is represented by *(p<0.05), **(p<0.01), ***(p<0.001) and ****(p<0.0001).

In all BrM samples, bone-marrow derived macrophages were the major tumor infiltrating immune cell type, comprising 3% of all DAPI^+^ cells in tumors and present at a median density of 130 cells/mm^2^, followed by CD3^+^ T-cells which comprised 1% of DAPI^+^ cells and were present at a density of 27 cells/mm^2^ (Figure 2B and Figure S2A). Dendritic cells and microglia were the least abundant immune cell type, comprising less than 1% of cells in the tumor tissue examined (Figure 2B). Similar macrophage and T-cell abundance and density was observed in metastatic brain tumors based on primary sites - breast and lung (Figure 2C-D and Figure S2D-E), and skin and thyroid (Figure S2B-C and Figure S2F-G). When considering the total immune cells analyzed, CD68^+^ macrophages comprise the biggest portion, followed by CD3^+^ T-cells, CD11c^+^ dendritic cells and TMEM119^+^ microglia across lung, breast, skin and thyroid BrM (Figure 2E).

### Macrophages and T-cells preferentially localize to ECM-rich regions

As we observed extensive collagen deposition in BrM tissue (Figure 1), we then examined whether the collagen deposition and organization impacted immune cell composition and localization within ECM-rich and neoplastic cell (NC)-rich regions. Hematoxylin and eosin and Masson’s Trichrome staining were used to annotate and distinguish between ECM-rich and NC-rich regions (Figure 3A). Spatial analysis indicated that a larger proportion of CD68^+^ macrophages were localized within NC-rich regions, compared to the ECM-rich regions in BrM (Figure 3B). The data also demonstrated that while CD68^+^ macrophages were more abundant in NC-rich regions, macrophage cell density was higher in the ECM-rich region compared to the NC-rich region (Figure 3C). The proportion of CD3^+^ T-cells was not statistically different between NC and ECM-rich regions (Figure 3D). However, T-cell density was significantly higher in ECM-rich areas compared to NC-rich areas (Figure 3E). A similar trend was observed in the distribution of CD68+ macrophages and CD3^+^ T-cells across BrM of primary tumors from the lung, breast (Figure 3F-I) and thyroid but not skin (Figure S3E-H). The data also shows an equal distribution of CD11c^+^ dendritic cells and TMEM119^+^ microglia across the entire tissue, with no significant differences in cell density of these cell types between NC- and ECM-rich regions in all BrM tissue cores analyzed (Figure S3A-D) and individual BrM tumor types (Figure SI-L). To determine the distribution of macrophages and T-cells in BrM tissue, infiltration analysis was performed to measure cell distribution in 50μm wide zones across the NC-rich tissue, NC-ECM interface, and within the ECM-rich compartment (Figure 4A). We observed that macrophages occupied two major regions. They were observed within ECM-rich regions and within NC-rich zones, infiltrating 100μm into the NC-rich area, with the highest macrophage NC-rich infiltration observed in lung BrM (Figure 4B). By contrast, most T-cells were localized in ECM-rich regions and did infiltrate further than 50μm into the NC-rich region, suggesting that T-cells do not infiltrate deep into the tumor in lung BrM, compared to macrophages (Figure 4D). A similar macrophage and T-cell distribution was observed in breast BrM tissue (Figure 4C and 4E).

**Figure 3.**
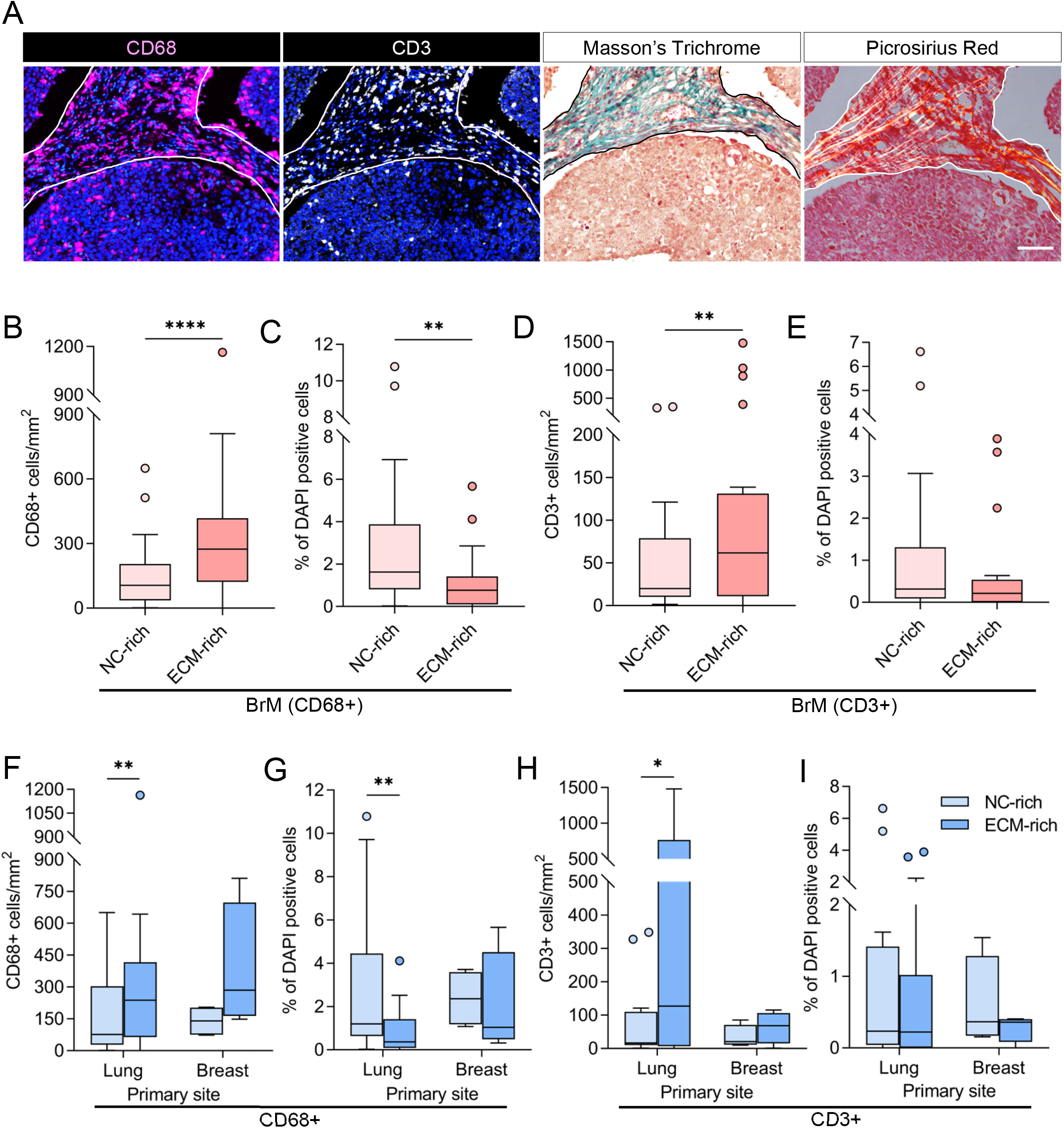
Preferential localization of T-cells and macrophages in ECM-rich regions. (**A**) Representative mIHC images of CD68^+^ and CD3^+^ cells localized in collagen-rich regions, Masson’s Trichrome and Picrosirius red staining of the same tissue region. Scale bars, 100μm. (**B**) CD68^+^ cell density (cells/mm^2^) and (**C**) CD68^+^ cells as a proportion of total DAPI^+^ cells in NC- and ECM-rich BrM tissue (n=24) regions, identified by Masson’s Trichrome staining. (**D**) CD3^+^ cell density (cells/mm^2^) and (**E**) CD3^+^ cells as a proportion of total DAPI^+^ cells (whole tissue core) in NC- and ECM-rich BrM tissue (n=22) regions identified by Masson’s Trichrome staining. (**F**) Cell density of CD68^+^ (cells/mm^2^) and (**G**) proportion of CD68^+^ cells of total DAPI^+^ cells in NC- and ECM-rich regions in BrM from lung (n=14) and breast tissue (n=4). (**H**) Cell density of CD68^+^ (cells/mm^2^) and (**I**) the proportion of CD68^+^ cells of total DAPI^+^ cells in NC- and ECM-rich regions in lung and breast BrM. All results are presented as median +/− IQR. Statistical significance was determined by Wilcoxon matched-pair signed rank test. Significance is represented by *(p<0.05), **(p<0.01), ***(p<0.001) and ****(p<0.0001).

**Figure 4.**
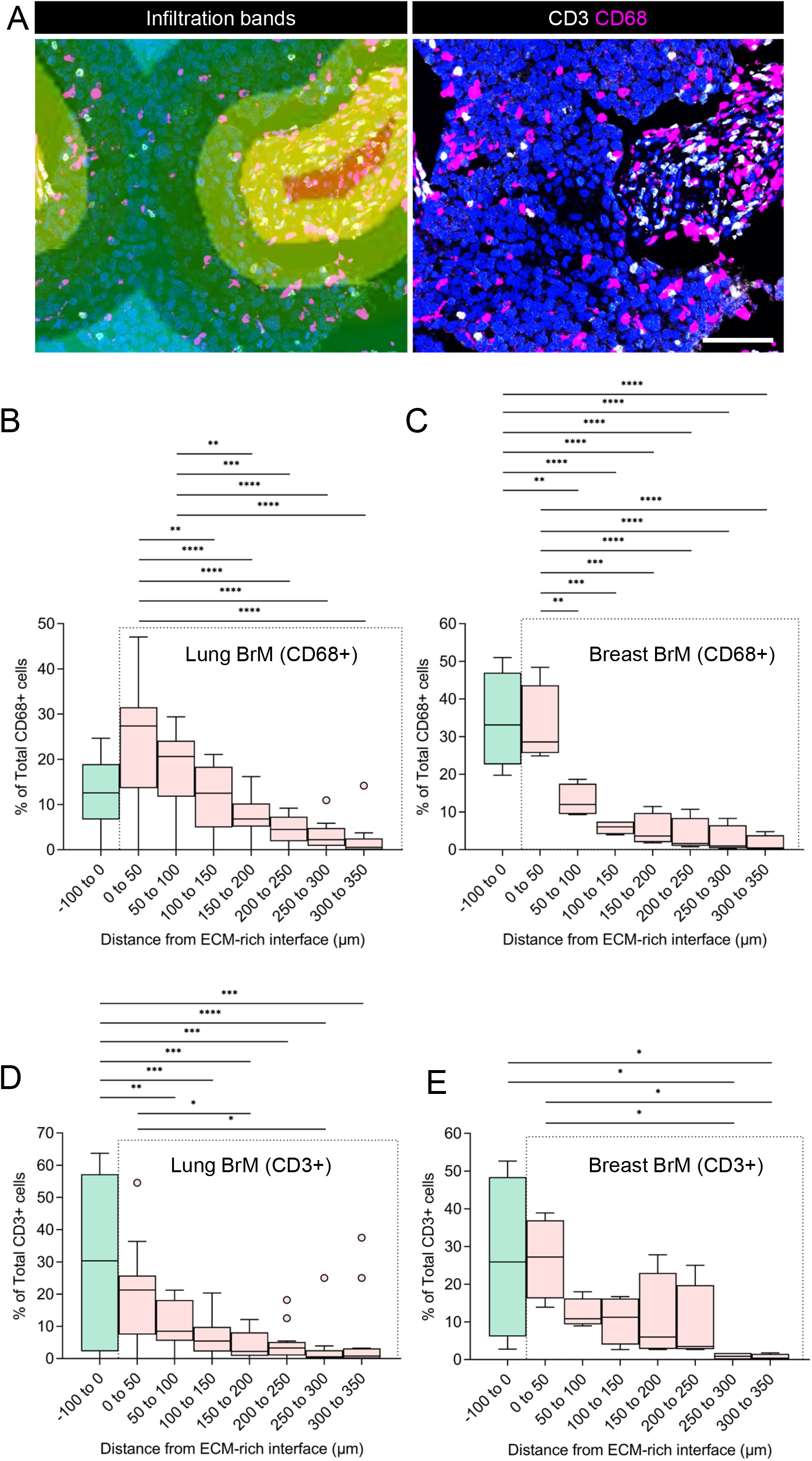
CD3^+^ T-cell and CD68^+^ macrophage infiltration in BrM. (**A**) Representative images showing infiltration bands/zones (upper panel), used to measure the number of CD68^+^ and CD3^+^ cells at different distances. (**B-C**) CD68^+^ and (**D-E**) CD3^+^ cell localization, as the distance from the NC-rich and ECM-rich interface, was determined (as a percentage of total C68^+^ cells or CD3^+^ cells) and expressed as an infiltration distance from ECM-rich regions (−100 to 0μm) into NC-rich regions (0 to 350μm). Data is shown for BrM from lung (**B and D**) (n=12) and breast (**C and E**) (n=4) BrM tumors. Statistical significance was determined by an ordinary one-way ANOVA. Significance is represented by *(p<0.05), **(p<0.01), ***(p<0.001) and ****(p<0.0001).

### Most macrophages are immunosuppressive and localize to ECM-rich stromal regions

To determine the proportion of macrophage subsets in BrM, serial tissue sections were analyzed by mIHC, using a myeloid cell-specific antibody panel to identify pro- and anti-inflammatory macrophage subsets. Antibodies recognizing Human Leukocyte Antigen-DR isotype (HLA-DR), CD163, CD206, CD68 TMEM119 and AE1/AE3 were used (Figure 5A). There were at least six different macrophage subsets, identified by the co-expression of pro- and anti-inflammatory biomarkers (Figure 5B, Table 1).

**Figure 5.**
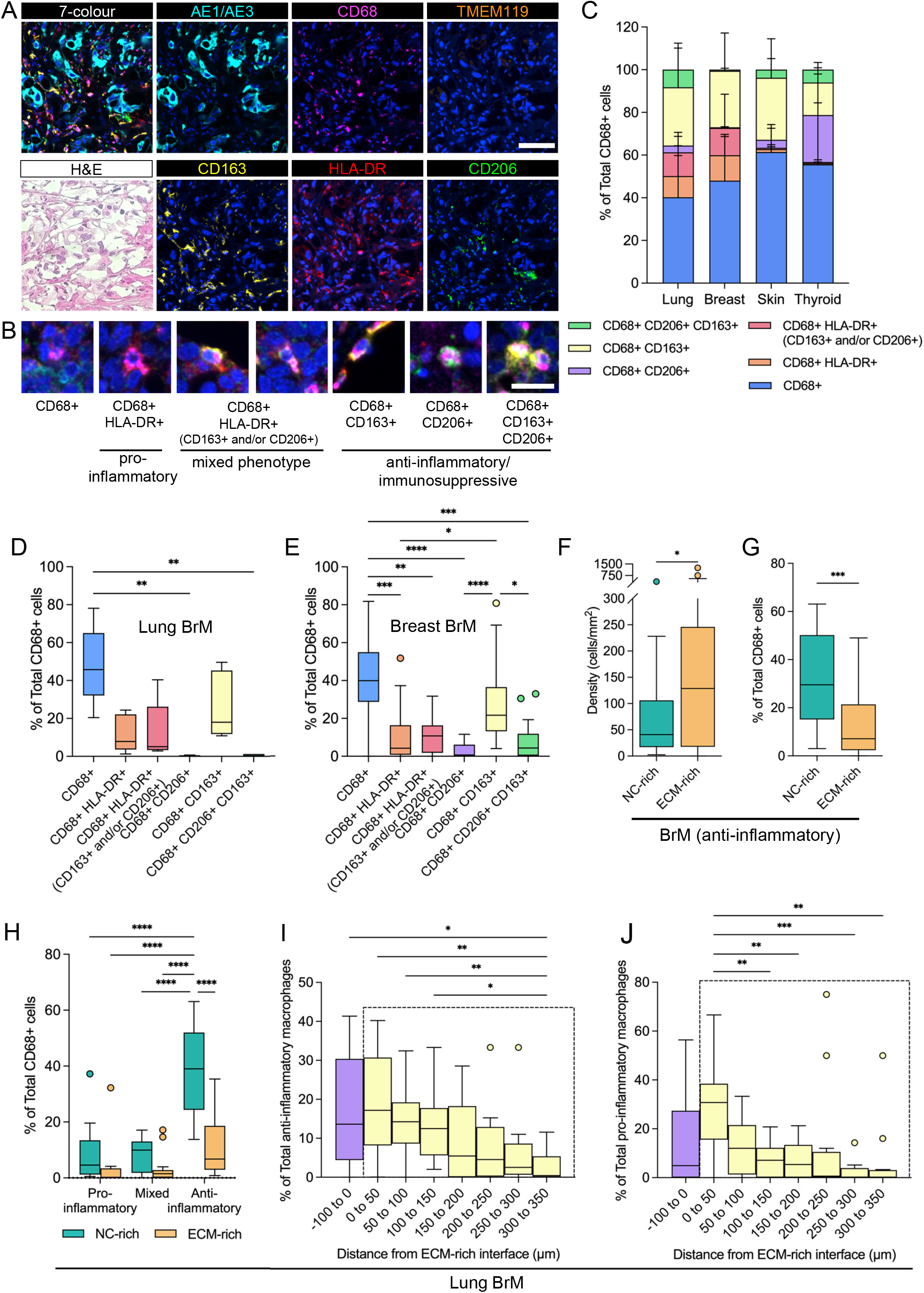
Macrophage subset localization in BrM. (**A**) Representative images of H&E and mIHC staining for AE1/AE3, CD68, CD163, CD206, HLA-DR and TMEM119 antibodies. Scale bars, 100μm. (**B**) Representative images showing the pro- and anti-inflammatory, and mixed phenotype of macrophages based on HLA-DR, CD163 and CD206 antibody staining. Scale bars, 25μm. (**C**) The proportion of different macrophage subsets, as a proportion of total CD68+ cells in BrM from lung (n=18), breast (n=5), skin (n=3) and thyroid (n=3). Results are presented as mean +/− SD. Percentage of macrophage subsets (CD68^+^, CD68^+^ HLA-DR^+^, CD68^+^ HLA-DR^+^ CD163^+^ and/or CD206^+^, CD68^+^ CD163^+^, and CD68^+^ CD206^+^ CD163^+^) as a proportion of total CD68^+^ cells in (**D**) lung (n=18) and (**E**) breast BrM (n=5) tissue. Statistical significance was determined by Kruskal-Wallis test, followed by Dunn’s test for multiple comparisons (α=0.05). (**F**) Percentage of anti-inflammatory macrophages subsets as a proportion of total CD68^+^ cells (whole tissue core) in NC-rich and ECM-rich regions (n=24) of BrM tissue. (**G**) Cell density (cells/mm^2^) of anti-inflammatory macrophages in NC-rich and ECM-rich regions (n=24). (**H**) Pro-inflammatory, anti-inflammatory and mixed subset macrophages, as a proportion of total CD68+ cells (whole tissue core) in NC-rich and ECM-rich regions in lung BrM tissue (n=14). Statistical significance was determined by Wilcoxon matched-pairs signed rank test. (**I**) The percentage of anti-inflammatory and (**J**) pro-inflammatory macrophage subset cell localization shown as the distance from the NC-rich and ECM-rich interface (zero distance point). The X-axis shows the infiltration distance within the ECM-rich region (negative values, −100 to 0μm) and into the NC-rich region (positive values, 0 to 350μm, within the dotted box). Data is shown for lung BrM tissue (n=14). All results are presented as median +/− IQR. Statistical significance was determined by an ordinary one-way ANOVA. Significance is represented by *(p<0.05), **(p<0.01), ***(p<0.001) and ****(p<0.0001).

The data shows that the predominant macrophage subset in all BrM types was the single positive CD68^+^, comprising between 40 and 60% of all CD68^+^ cells, followed by the anti-inflammatory CD68^+^CD163^+^ subset, comprising 20-30% of CD68^+^ cells (Figure 5C). The pro-inflammatory CD68^+^ HLA-DR^+^ subset comprised about 10% of macrophages in lung and breast BrM, but not in skin and thyroid BrM (Figure 5D-E, Figure S4A-B). Moreover, a subset of macrophages co-expressing HLA-DR and CD163, and/or CD206 was identified (Table 1, Figure 5B). This mixed phenotype constituted approximately 10% of macrophages in lung and breast BrM (Figure 5D-E). Our results demonstrate that different macrophages subsets are present in BrM and the proportion differs depending on the primary tumor type.

Using CD163 and CD206 expression in macrophages as a measure of the anti-inflammatory (immunosuppressive) macrophage subset, the total density and proportion of CD68^+^CD206^+^, CD68^+^CD163^+^, and CD68^+^CD163^+^CD206^+^ cells in ECM-rich and NC-rich regions was measured in all BrM samples (Figure 5F-G). There was a higher density of immunosuppressive macrophages in ECM-rich regions, at a median density of 129 cells/mm^2^, compared to the NC-rich, with a median cell density of 41 cells/mm^2^ (Figure 5F). By contrast, the proportion of this macrophage subset relative to total CD68^+^ cells was higher in NC-rich regions, at 29.5%, compared to ECM-rich regions, at 7.1% (Figure 5G). Next, we compared the proportion of these anti-inflammatory cells with pro-inflammatory (CD68^+^HLA-DR^+^) macrophages and macrophages with a mixed phenotype (CD68^+^ HLA-DR^+^ (CD163^+^ and/or CD206^+^)) (Figure 5H). Anti-inflammatory macrophages were present in a higher proportion than other subsets, at approximately 20 - 50%, in NC-rich regions in BrM from lung, breast and thyroid primary tumors (Figure 5H, Figure S4C and S4E). Interestingly, skin cancer BrM showed a higher proportion of anti-inflammatory macrophages in ECM-rich regions (Figure S4D). Overall, we observed the predominance of anti-inflammatory macrophages within ECM-rich regions in BrM tissue, compared with pro-inflammatory and other macrophage subsets.

We also investigated the distribution of pro- and anti-inflammatory macrophages within NC-rich areas. As described in the previous section, we performed infiltration analysis to determine cell localization in ECM-rich regions and macrophage localization, expressed as the distance from NC-rich regions. In lung BrM samples, we observed an equal distribution of anti-inflammatory cells (median 15% of total anti-inflammatory macrophages) in 0-50μm, 50-100μm, and 100-150μm bands in the NC-rich regions (Figure 5I). By contrast, 30% of pro-inflammatory and mixed macrophages were within 50μm of NC-rich regions (Figure 5J and Figure S5A). A similar distribution pattern of anti-, pro-inflammatory and mixed macrophages in the NC-rich regions was also observed in metastatic tumors from breast, skin and thyroid primaries (Figure S5B-J). The data suggest that anti-inflammatory macrophages are distributed more evenly across the NC-rich regions compared to pro-inflammatory and mixed macrophages.

### Macrophage and T-cell abundance correlates with collagen content in BrM

Given the differences between immune cell localization between ECM-rich or NC-rich regions in BRM, we investigated whether the presence of collagen correlates with the presence of immune cells. Correlation analysis revealed a strong correlation between collagen content and total CD68^+^ macrophages and CD3^+^ T-cell abundance (as a proportion of all DAPI^+^ cells and cell density) in BrM tumor tissue (Figure 6 and S6). Comparison between the various macrophage subsets and the presence of collagen, demonstrated a moderate correlation between the mixed-macrophage phenotype and collagen deposition. A strong correlation was also found between the pro- and mixed macrophage phenotypes, as well as between the presence of CD3^+^ T-cells and CD68^+^ macrophages (Figures 6 and S6). When this association between CD3^+^ T-cells and CD68^+^ macrophages was further assessed in terms of association with macrophage subset, we found that the presence CD3^+^ T-cells correlated with anti-inflammatory and mixed subset macrophages but not pro-inflammatory macrophages.

**Figure 6.**
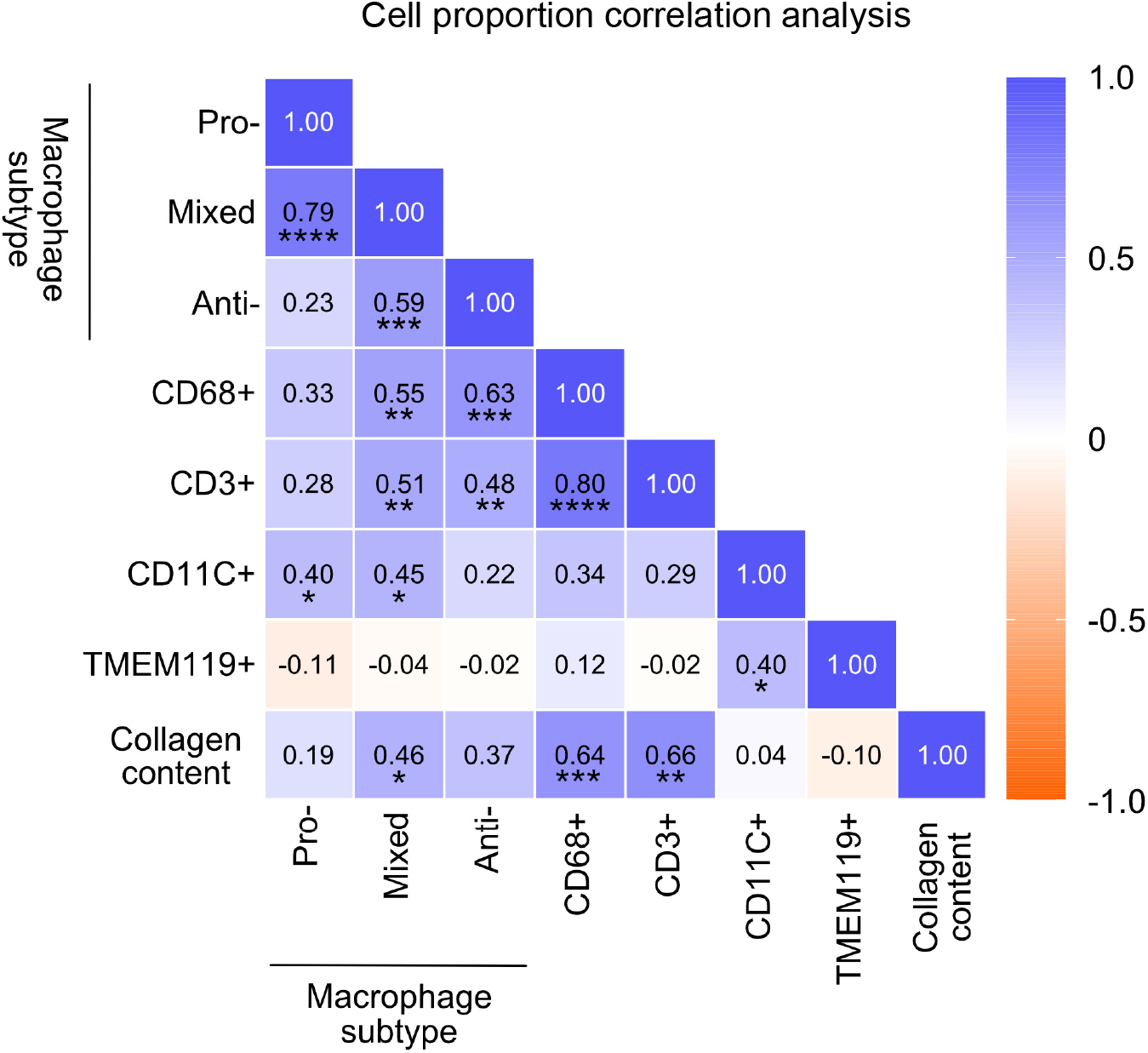
Correlation between macrophage number, T-cell number, and ECM content in BrM. Correlation heatmap between CD68^+^ (total CD68^+^), TMEM119^+^, CD11c^+^ and CD3^+^ cells, and pro-inflammatory, anti-inflammatory and mixed macrophage subsets (percentage of total DAPI cells) with collagen content in BrM tissue cores (n=28). Spearman rank correlation coefficients are indicated on the heatmap and significance is represented by *(p<0.05), **(p<0.01), ***(p<0.001) and ****(p<0.0001).

## Discussion

The complex network of extracellular proteins contributing to ECM remodeling underpins the structural organization of the tumor microenvironment and influences tumorigenicity by establishing niches which allow cancer cells to thrive. Collagen proteins are one of the major classes of ECM proteins in tumor tissue and the role of specific collagen proteins have been investigated in primary cancers. Metastatic spread of tumor cells from the primary site are associated with advanced disease and poor prognosis for patients with cancer. Metastatic tumors are the most common neoplasm affecting the brain [28]. Here, we demonstrate that brain metastatic tumor tissue exhibits substantial ECM deposition which compartmentalizes the stroma from the neoplastic cell rich tissue.

ECM content, measured as the proportion of collagen-stained area, in lung and breast cancer BrM tissue was similar to that measured in GBM tissue. However, the organization of the collagen-rich ECM stroma in BrM differed compared with GBM. All BrM tissues exhibited ECM-rich stromal regions with well-defined borders at the NC-rich region interface, and the NC-rich regions exhibited a mosaic-like appearance, bordered by interstitial-like ECM stroma. The collagen in BrM tissue was composed of thicker, denser collagen bundles, compared to GBM in which the collagen exhibited thinner, less dense fibrillar collagens. The highly organized appearance of BrM tumor tissue, compared with GBM tissue, could be explained, in part, by the nature of the metastatic cell type and the spectrum and level of ECM protein expression. For example, among the major twenty cancer types, collagen I mRNA and protein expression is lowest in GBM cells, while collagen IV expression levels in GBM is similar to other cancer types [29] [30]. Another contributing factor resulting in ECM content and organization between primary brain tumors such as GBM and BrM, is the infiltrative nature of GBM cells which result in diffuse collagen deposition in the brain parenchyma. These differences would result in distinct ECM protein expression, deposition patterns and collagen fiber maturation.

Tumor infiltrating macrophages were the major immune cell type detected in BrM tumors, irrespective of tumor origin. Infiltrating T-cells was the next most abundant immune cell type, while the brain resident myeloid cells, microglia were present at relatively low numbers in lung and skin BrMs and no detectable microglial infiltration in breast BrM. These observations demonstrate that specific mechanisms operate in the establishment and development of the host tissue immune microenvironment and depend on metastatic cell ontogeny. Previous studies investigating differences between tumor infiltrating macrophages and microglia in GBM and BrM did not report differences in immune cell properties and function, between GBM and BrM [31]. Cell number correlation analysis also showed a strong link between ECM and the number of immune cells, suggesting that immune cells preferentially localize to ECM-rich regions, or that immune cells may contribute to the ECM content in BrM. The correlation between T-cells and macrophages, suggests that the presence of one cell type acts as an attractant for the other cell type or that both cell types are attracted to the same factor(s). Overall, our data shows that metastatic brain tumors exhibit substantial ECM deposition, and that the collagen fiber property in the BrM ECM differs compared to the collagen in GBM. We also show that immune cells in BrM are preferentially localized to ECM-rich regions and that most bone marrow derived macrophages are immunosuppressive. This suggests that like GBM, tumor infiltrating macrophages are key players in establishing an immunosuppressive tumor microenvironment. The findings support the view that optimal therapies for patients with metastatic brain disease need to consider therapies which modify the disease-specific tumor matrisome and the targeting of immunosuppressive macrophages, in combination with current cytotoxic and/or cytostatic therapies.

## Acknowledgments

We thank Dr. Metta Jana and Dr. Rejhan Idrizi from Centre for Advanced Histology and Microscopy (Peter MacCallum Cancer Centre) and Dr. Ian Birchall (The Florey Institute of Neuroscience and Mental Health) for advice on tissue preparation, histology, and staining protocols. MD and MP were supported by the Australian Government Research Training Program Scholarship. SSW was supported by a University of Melbourne, Melbourne Research Scholarship. TRC was supported by the National Health and Medical Research Council (NHMRC) (1158590). SSS was supported by an MRFF Accelerated Research - Australian Brain Cancer Mission grant (1158175). TM was supported by a CASS Foundation Science Grant (065516), a Brain Foundation Grant (065518) and a Department of Surgery and School of Biomedical Sciences collaborative science grant (2018).

## Supplementary Data

**Table S1.**
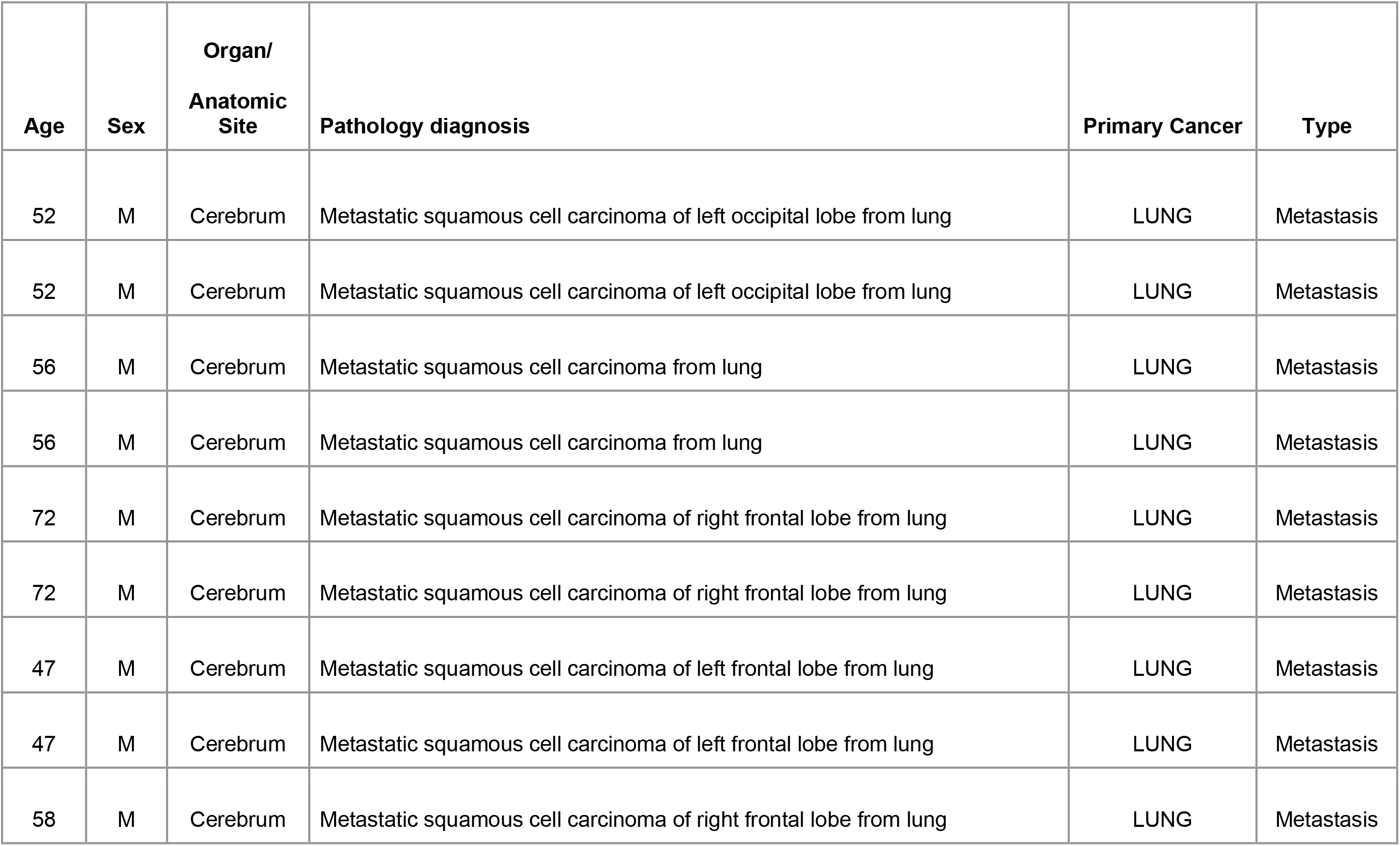
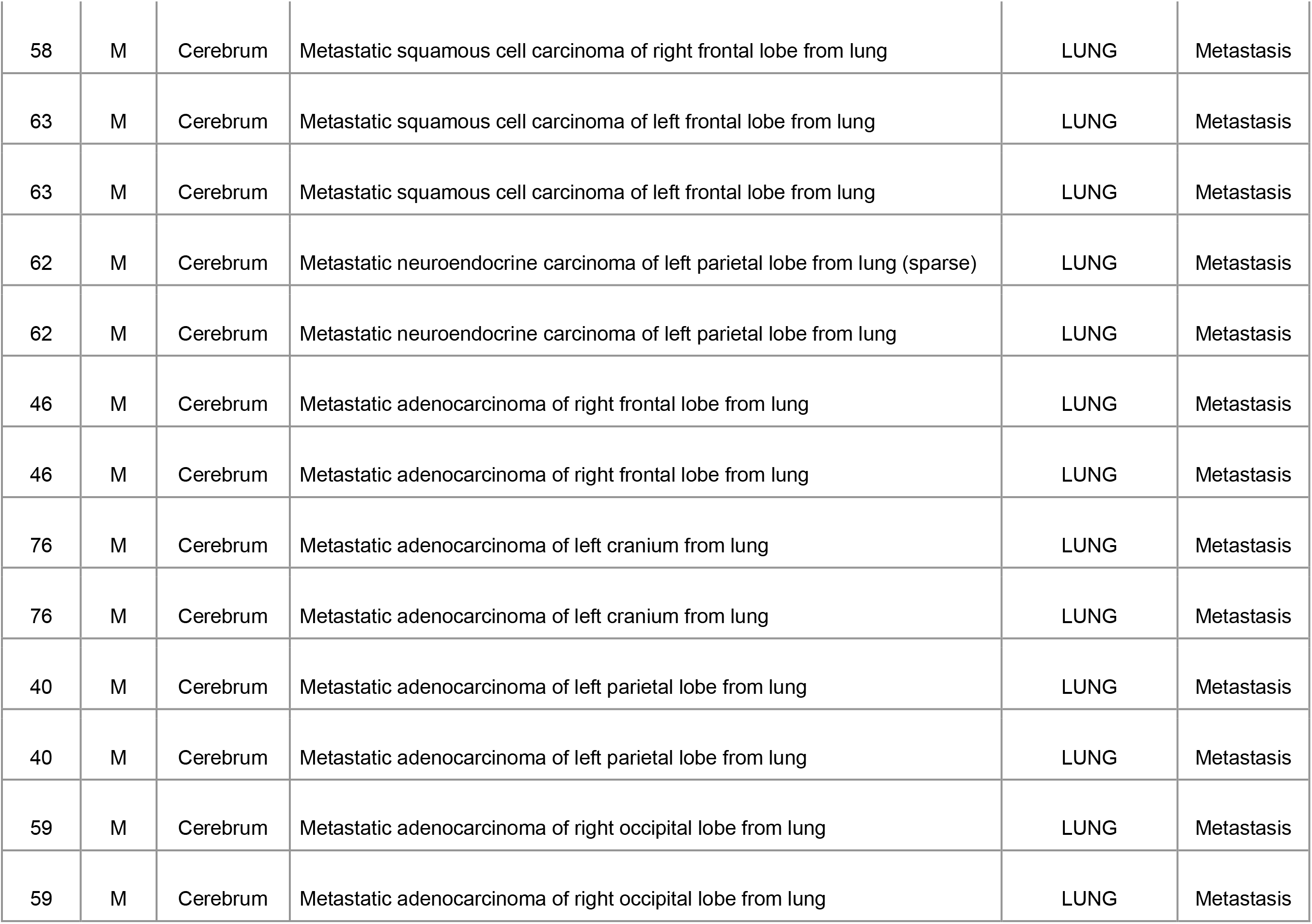
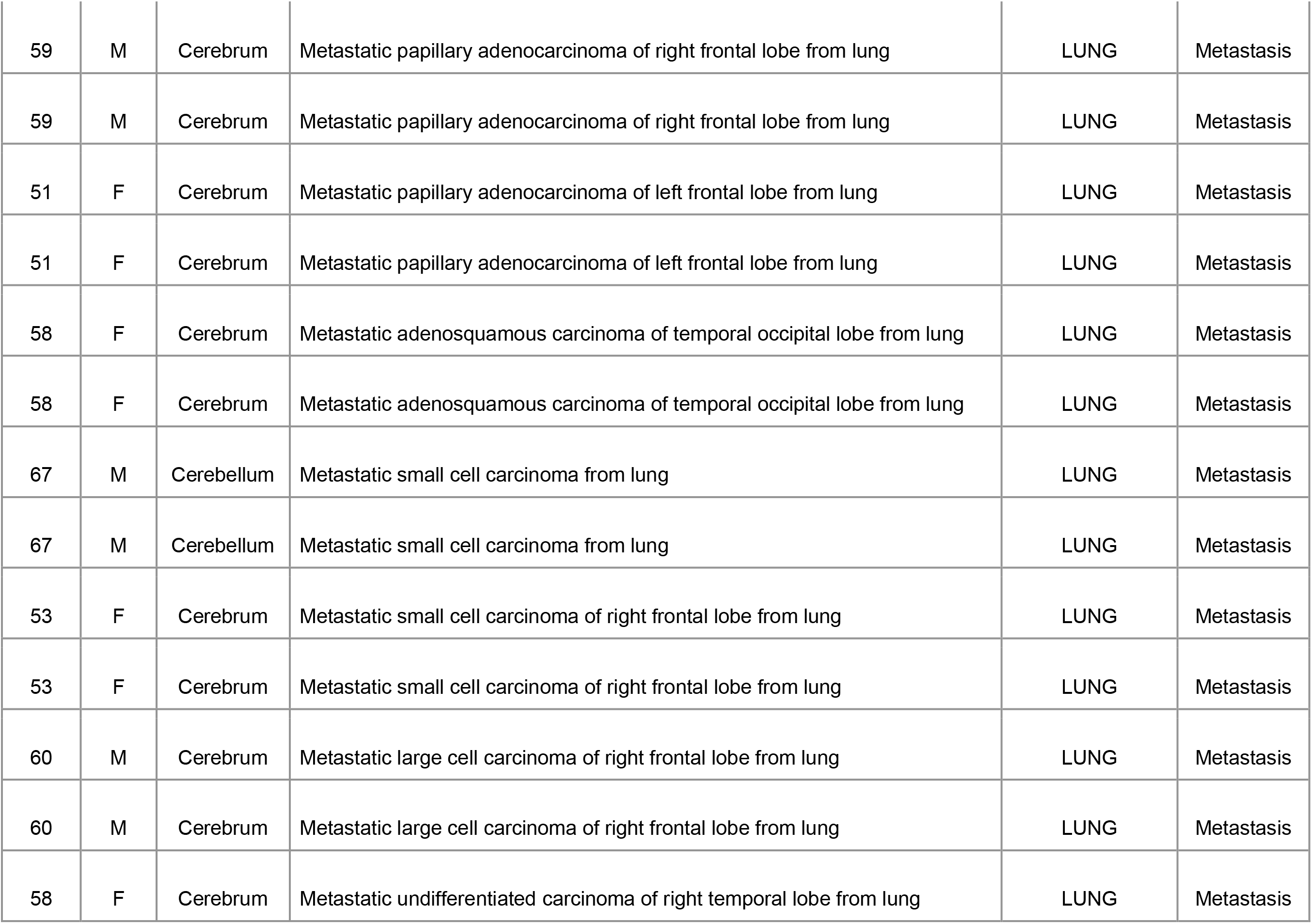
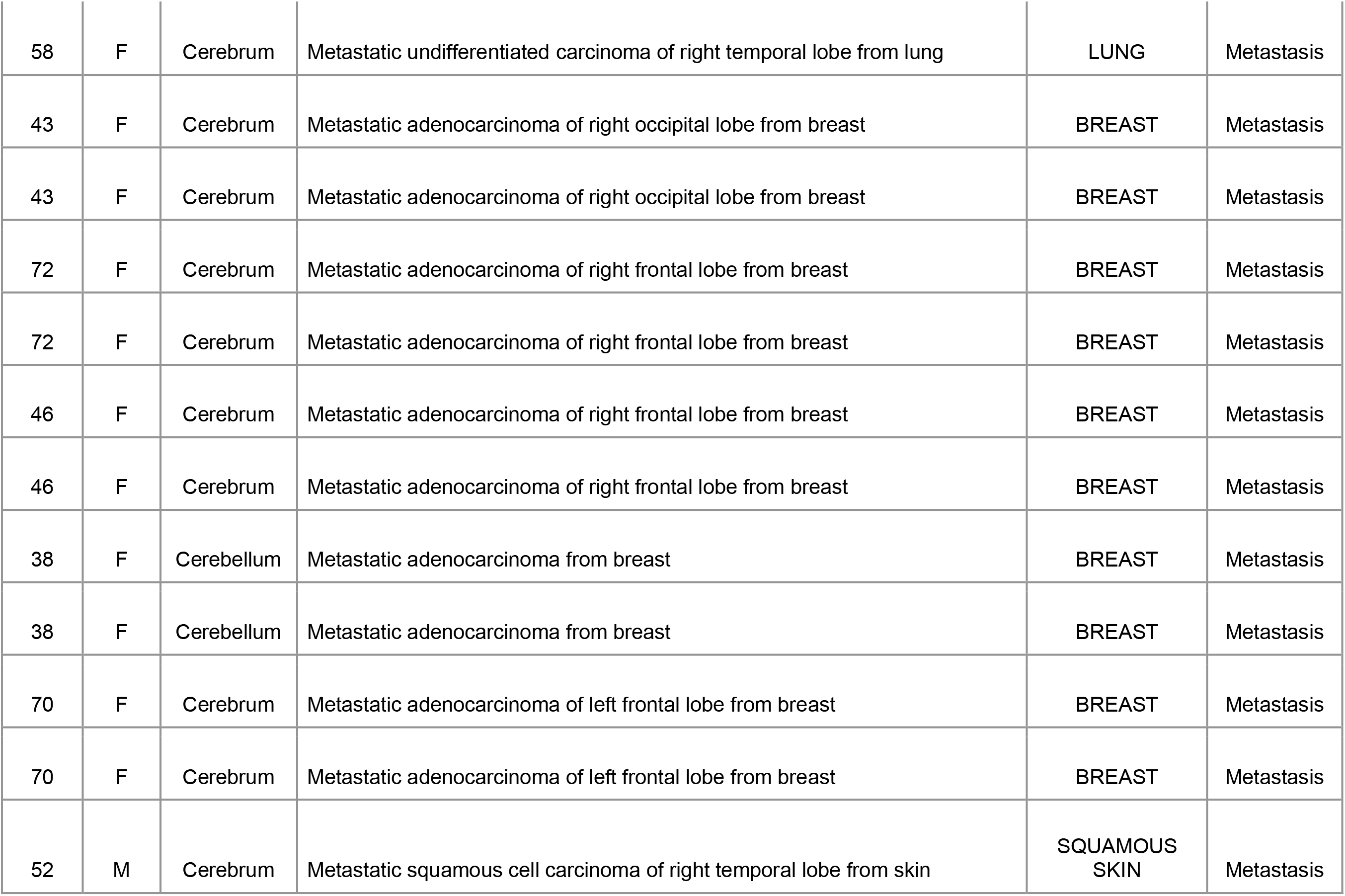
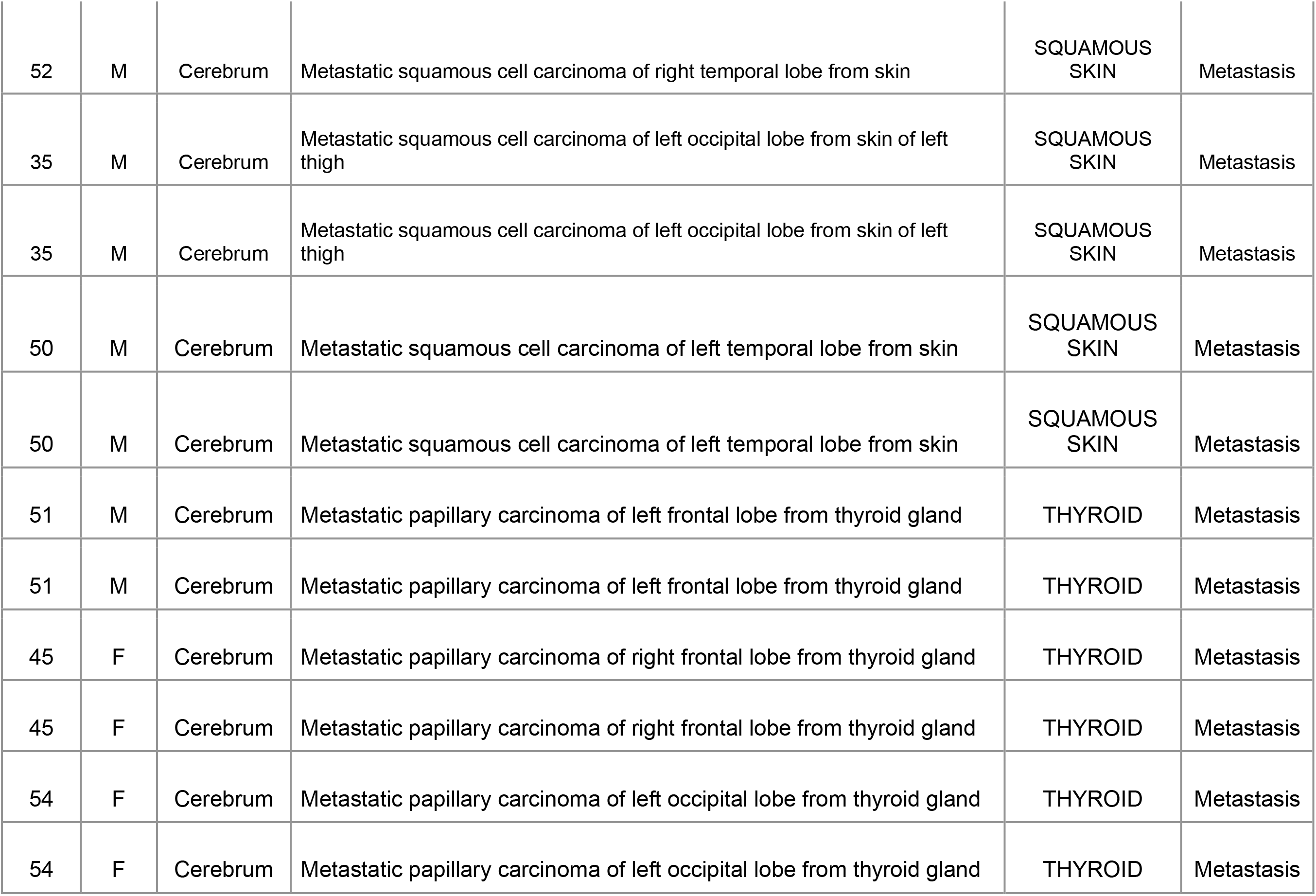

| **Age** | **Sex** | **Organ/**<br>**Anatomic**<br>**Site** | **Pathology diagnosis** | **Primary Cancer** | **Type** |
| --- | --- | --- | --- | --- | --- |
| 52 | M | Cerebrum | Metastatic squamous cell carcinoma of left occipital lobe from lung | LUNG | Metastasis |
| 52 | M | Cerebrum | Metastatic squamous cell carcinoma of left occipital lobe from lung | LUNG | Metastasis |
| 56 | M | Cerebrum | Metastatic squamous cell carcinoma from lung | LUNG | Metastasis |
| 56 | M | Cerebrum | Metastatic squamous cell carcinoma from lung | LUNG | Metastasis |
| 72 | M | Cerebrum | Metastatic squamous cell carcinoma of right frontal lobe from lung | LUNG | Metastasis |
| 72 | M | Cerebrum | Metastatic squamous cell carcinoma of right frontal lobe from lung | LUNG | Metastasis |
| 47 | M | Cerebrum | Metastatic squamous cell carcinoma of left frontal lobe from lung | LUNG | Metastasis |
| 47 | M | Cerebrum | Metastatic squamous cell carcinoma of left frontal lobe from lung | LUNG | Metastasis |
| 58 | M | Cerebrum | Metastatic squamous cell carcinoma of right frontal lobe from lung | LUNG | Metastasis |
| 58 | M | Cerebrum | Metastatic squamous cell carcinoma of right frontal lobe from lung | LUNG | Metastasis |
| 63 | M | Cerebrum | Metastatic squamous cell carcinoma of left frontal lobe from lung | LUNG | Metastasis |
| 63 | M | Cerebrum | Metastatic squamous cell carcinoma of left frontal lobe from lung | LUNG | Metastasis |
| 62 | M | Cerebrum | Metastatic neuroendocrine carcinoma of left parietal lobe from lung (sparse) | LUNG | Metastasis |
| 62 | M | Cerebrum | Metastatic neuroendocrine carcinoma of left parietal lobe from lung | LUNG | Metastasis |
| 46 | M | Cerebrum | Metastatic adenocarcinoma of right frontal lobe from lung | LUNG | Metastasis |
| 46 | M | Cerebrum | Metastatic adenocarcinoma of right frontal lobe from lung | LUNG | Metastasis |
| 76 | M | Cerebrum | Metastatic adenocarcinoma of left cranium from lung | LUNG | Metastasis |
| 76 | M | Cerebrum | Metastatic adenocarcinoma of left cranium from lung | LUNG | Metastasis |
| 40 | M | Cerebrum | Metastatic adenocarcinoma of left parietal lobe from lung | LUNG | Metastasis |
| 40 | M | Cerebrum | Metastatic adenocarcinoma of left parietal lobe from lung | LUNG | Metastasis |
| 59 | M | Cerebrum | Metastatic adenocarcinoma of right occipital lobe from lung | LUNG | Metastasis |
| 59 | M | Cerebrum | Metastatic adenocarcinoma of right occipital lobe from lung | LUNG | Metastasis |
| 59 | M | Cerebrum | Metastatic papillary adenocarcinoma of right frontal lobe from lung | LUNG | Metastasis |
| 59 | M | Cerebrum | Metastatic papillary adenocarcinoma of right frontal lobe from lung | LUNG | Metastasis |
| 51 | F | Cerebrum | Metastatic papillary adenocarcinoma of left frontal lobe from lung | LUNG | Metastasis |
| 51 | F | Cerebrum | Metastatic papillary adenocarcinoma of left frontal lobe from lung | LUNG | Metastasis |
| 58 | F | Cerebrum | Metastatic adenosquamous carcinoma of temporal occipital lobe from lung | LUNG | Metastasis |
| 58 | F | Cerebrum | Metastatic adenosquamous carcinoma of temporal occipital lobe from lung | LUNG | Metastasis |
| 67 | M | Cerebellum | Metastatic small cell carcinoma from lung | LUNG | Metastasis |
| 67 | M | Cerebellum | Metastatic small cell carcinoma from lung | LUNG | Metastasis |
| 53 | F | Cerebrum | Metastatic small cell carcinoma of right frontal lobe from lung | LUNG | Metastasis |
| 53 | F | Cerebrum | Metastatic small cell carcinoma of right frontal lobe from lung | LUNG | Metastasis |
| 60 | M | Cerebrum | Metastatic large cell carcinoma of right frontal lobe from lung | LUNG | Metastasis |
| 60 | M | Cerebrum | Metastatic large cell carcinoma of right frontal lobe from lung | LUNG | Metastasis |
| 58 | F | Cerebrum | Metastatic undifferentiated carcinoma of right temporal lobe from lung | LUNG | Metastasis |
| 58 | F | Cerebrum | Metastatic undifferentiated carcinoma of right temporal lobe from lung | LUNG | Metastasis |
| 43 | F | Cerebrum | Metastatic adenocarcinoma of right occipital lobe from breast | BREAST | Metastasis |
| 43 | F | Cerebrum | Metastatic adenocarcinoma of right occipital lobe from breast | BREAST | Metastasis |
| 72 | F | Cerebrum | Metastatic adenocarcinoma of right frontal lobe from breast | BREAST | Metastasis |
| 72 | F | Cerebrum | Metastatic adenocarcinoma of right frontal lobe from breast | BREAST | Metastasis |
| 46 | F | Cerebrum | Metastatic adenocarcinoma of right frontal lobe from breast | BREAST | Metastasis |
| 46 | F | Cerebrum | Metastatic adenocarcinoma of right frontal lobe from breast | BREAST | Metastasis |
| 38 | F | Cerebellum | Metastatic adenocarcinoma from breast | BREAST | Metastasis |
| 38 | F | Cerebellum | Metastatic adenocarcinoma from breast | BREAST | Metastasis |
| 70 | F | Cerebrum | Metastatic adenocarcinoma of left frontal lobe from breast | BREAST | Metastasis |
| 70 | F | Cerebrum | Metastatic adenocarcinoma of left frontal lobe from breast | BREAST | Metastasis |
| 52 | M | Cerebrum | Metastatic squamous cell carcinoma of right temporal lobe from skin | SQUAMOUS<br>SKIN | Metastasis |
| 52 | M | Cerebrum | Metastatic squamous cell carcinoma of right temporal lobe from skin | SQUAMOUS SKIN | Metastasis |
| 35 | M | Cerebrum | Metastatic squamous cell carcinoma of left occipital lobe from skin of left thigh | SQUAMOUS SKIN | Metastasis |
| 35 | M | Cerebrum | Metastatic squamous cell carcinoma of left occipital lobe from skin of left thigh | SQUAMOUS SKIN | Metastasis |
| 50 | M | Cerebrum | Metastatic squamous cell carcinoma of left temporal lobe from skin | SQUAMOUS SKIN | Metastasis |
| 50 | M | Cerebrum | Metastatic squamous cell carcinoma of left temporal lobe from skin | SQUAMOUS SKIN | Metastasis |
| 51 | M | Cerebrum | Metastatic papillary carcinoma of left frontal lobe from thyroid gland | THYROID | Metastasis |
| 51 | M | Cerebrum | Metastatic papillary carcinoma of left frontal lobe from thyroid gland | THYROID | Metastasis |
| 45 | F | Cerebrum | Metastatic papillary carcinoma of right frontal lobe from thyroid gland | THYROID | Metastasis |
| 45 | F | Cerebrum | Metastatic papillary carcinoma of right frontal lobe from thyroid gland | THYROID | Metastasis |
| 54 | F | Cerebrum | Metastatic papillary carcinoma of left occipital lobe from thyroid gland | THYROID | Metastasis |
| 54 | F | Cerebrum | Metastatic papillary carcinoma of left occipital lobe from thyroid gland | THYROID | Metastasis |

**Figure S1.**
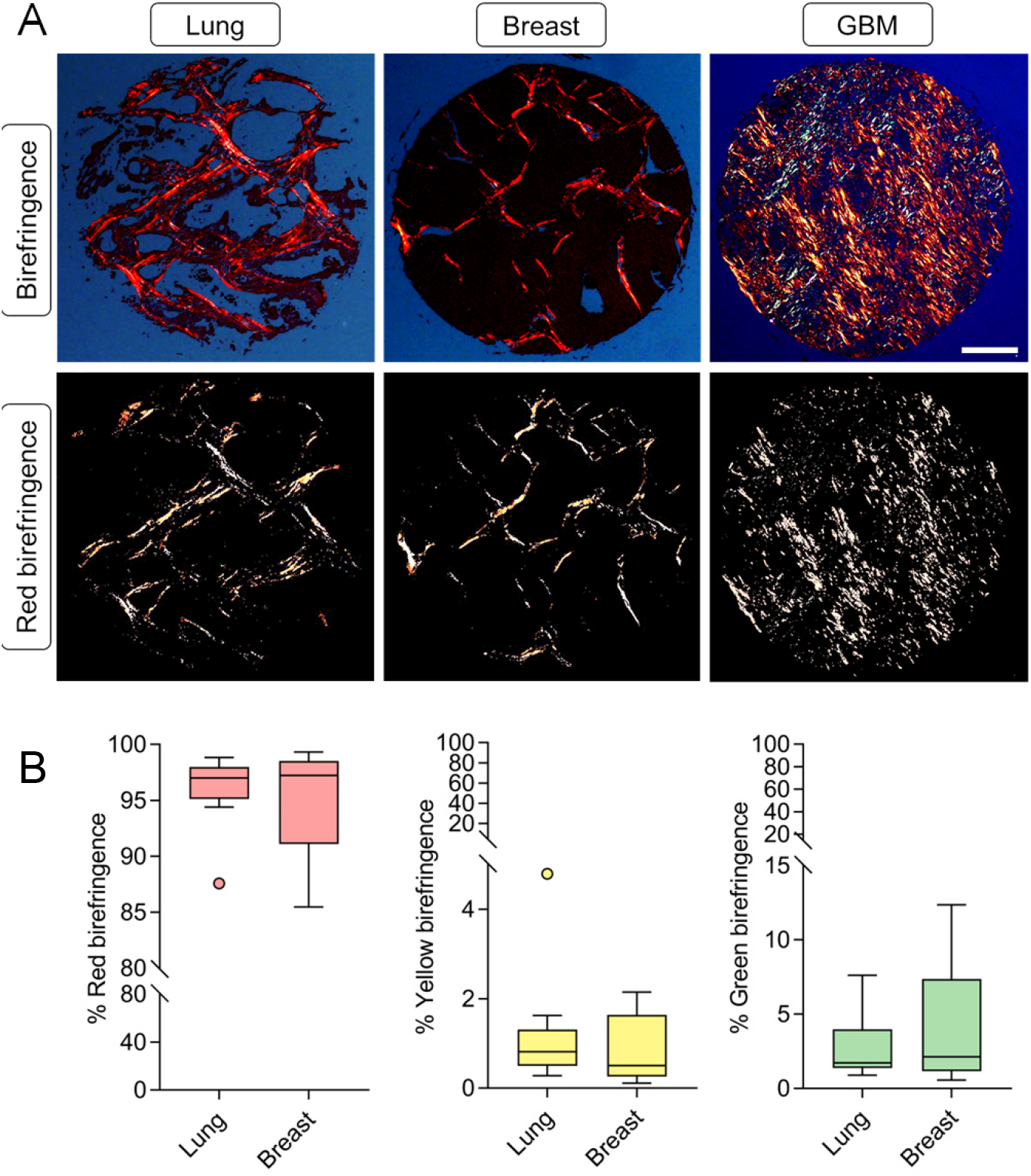
Collagen density and birefringence in lung and breast BrM. (**A**) Representative images of birefringence (polarized light) and red birefringence signal of BrM and GBM tumor tissue. Scale bar, 300μm. (**B**) Percentage of red, yellow and green birefringence signal of lung BrM (n=9) and breast BrM (n=5) tissue cores. Statistical significance of the birefringence signal (red, yellow and green) was determined by a Mann-Whitney test.

**Figure S2.**
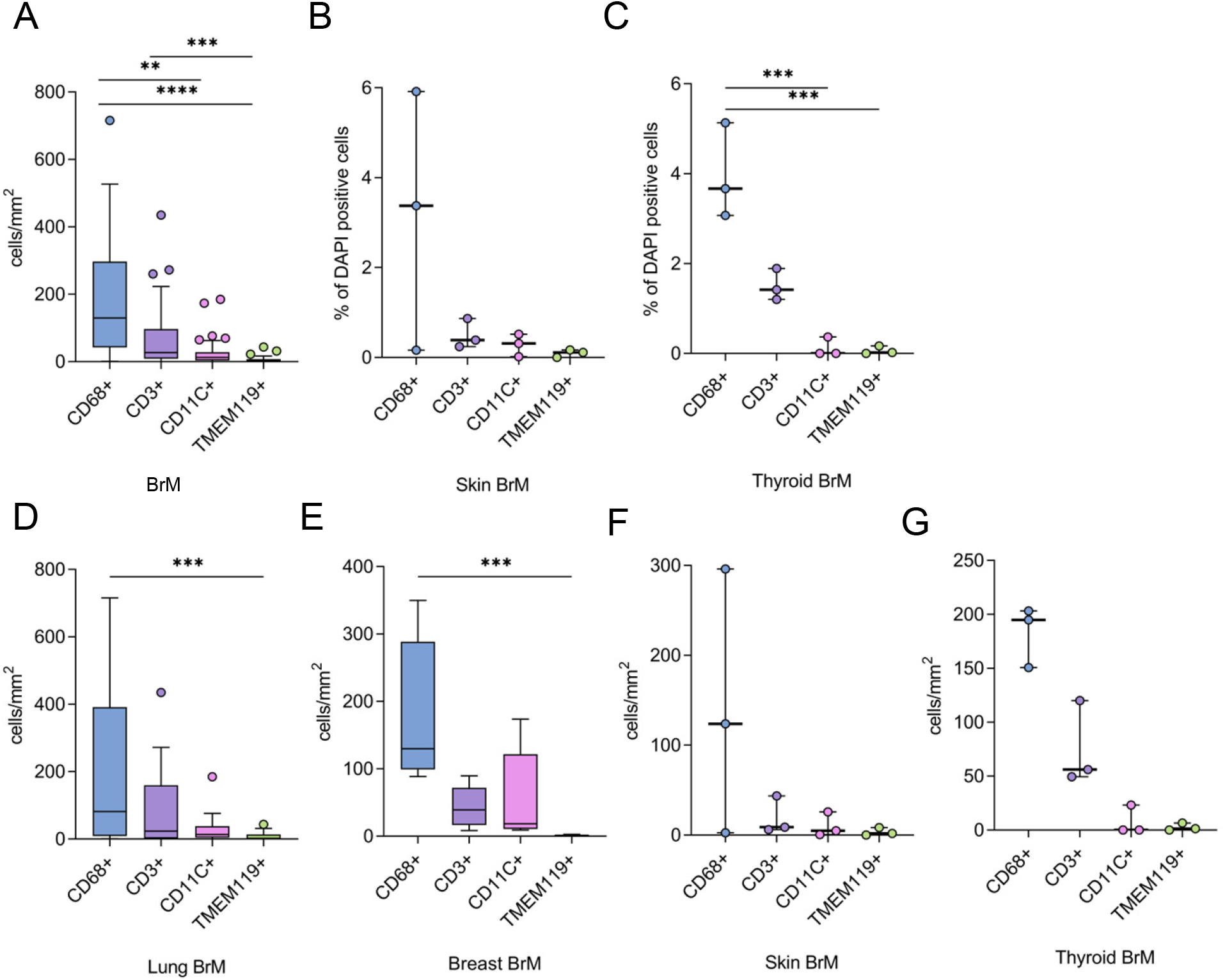
Macrophages are the major infiltrating immune cell type in BrM. (**A**) CD68^+^, CD11c^+^, TMEM119^+^ and CD3^+^ immune cell density (cells/mm^2^) in all BrM tumor tissue cores (n=29). Proportion of CD68^+^, CD11c^+^, TMEM119^+^ and CD3^+^ as a percentage of total DAPI^+^ cells in (**B**) skin (n=3) and (**C**) thyroid (n=3) BrM tissue. CD68^+^, CD11c^+^, TMEM119^+^ and CD3^+^ immune cell density (cells/mm^2^) in (**D**) lung (n=18), (**E**) breast (n=5), (**F**) skin (n=3) and (**G**) thyroid (n=3). Statistical significance was determined by Kruskal-Wallis test, followed by Dunn’s test for multiple comparisons (α=0.05) for comparison between immune cell types. Significance is represented by *(p<0.05), **(p<0.01), ***(p<0.001) and ****(p<0.0001).

**Figure S3.**
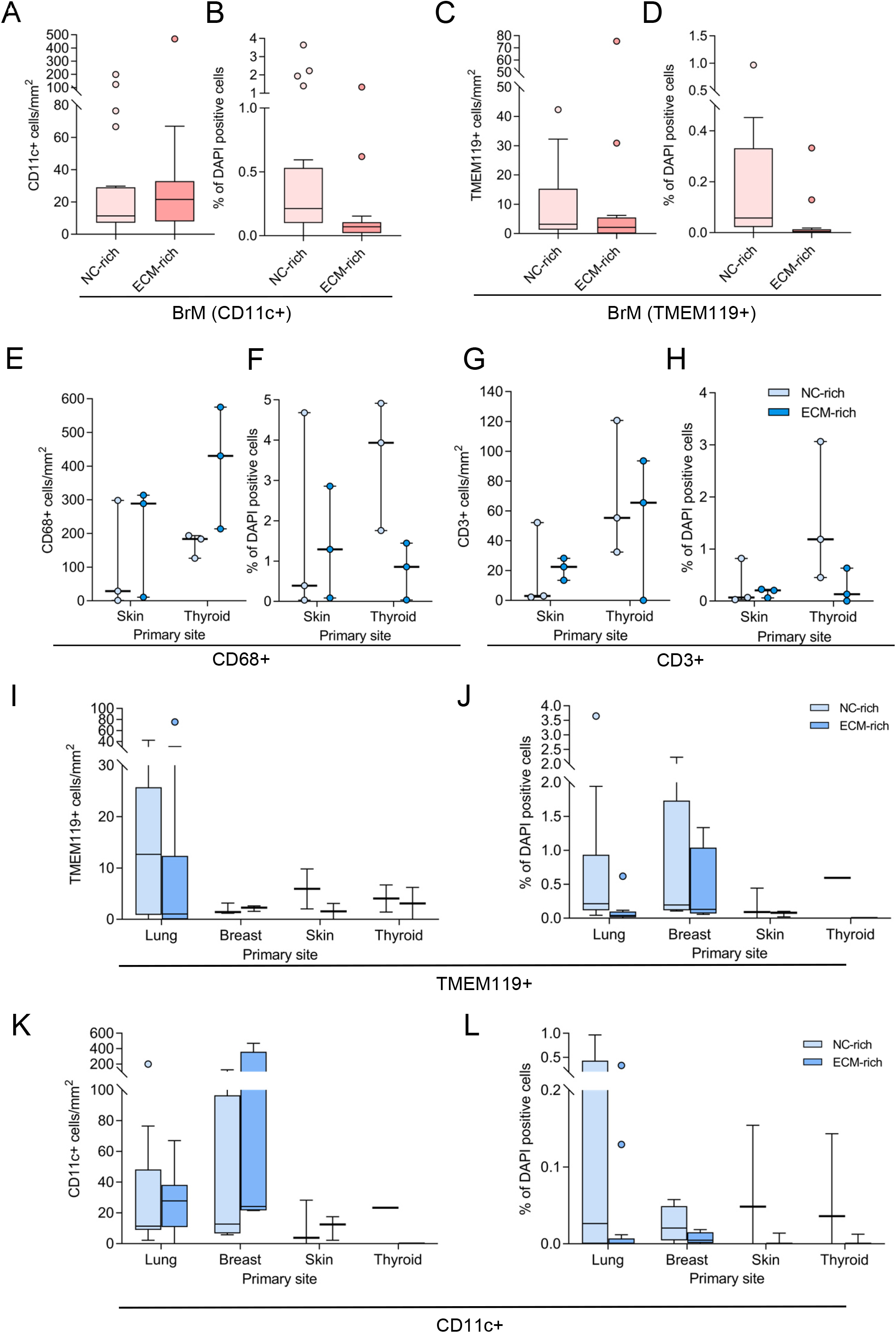
Preferential localization of T-cells and macrophages in ECM-rich regions. (**A**) CD11c^+^ cell density (cells/mm^2^) and (**B**) CD11c^+^ cells as a proportion of total DAPI^+^ cells (whole tissue core) in NC- and ECM-rich BrM tissue (n=21) regions. (**C**) TMEM119^+^ cell density (cells/mm^2^) and (**D**) TMEM119^+^ cells as a proportion of total DAPI^+^ cells (whole tissue core) in NC- and ECM-rich BrM tissue (n=17) regions. (**E**) CD68^+^ cell density (cells/mm^2^) and (**F**) CD68^+^ cells as a proportion of total DAPI^+^ cells (whole tissue core) in NC- and ECM-rich skin (n=3) and thyroid (n=3) BrM tissue regions. (**G**) CD3^+^ cell density (cells/mm^2^) and (H) CD3^+^ cells as a proportion of total DAPI^+^ cells (whole tissue core) in NC- and ECM-rich skin (n=3) and thyroid (n=3) BrM tissue regions. (**I**) TMEM119^+^ cell density (cells/mm^2^) and (**J**) TMEM119^+^ cells as a proportion of total DAPI^+^ cells (whole tissue core) in NC- and ECM-rich lung (n=14), breast (n=4), skin (n=3) and thyroid (n=3) BrM tissue. (**K**) CD11c^+^ cell density (cells/mm^2^) and (**L**) CD11c^+^ cells as a proportion of total DAPI^+^ cells (whole tissue core) in NC- and ECM-rich lung (n=13), breast (n=4), skin (n=3) and thyroid (n=1) BrM tissue. Statistical significance was determined by Wilcoxon matched-pairs signed rank test.

**Figure S4.**
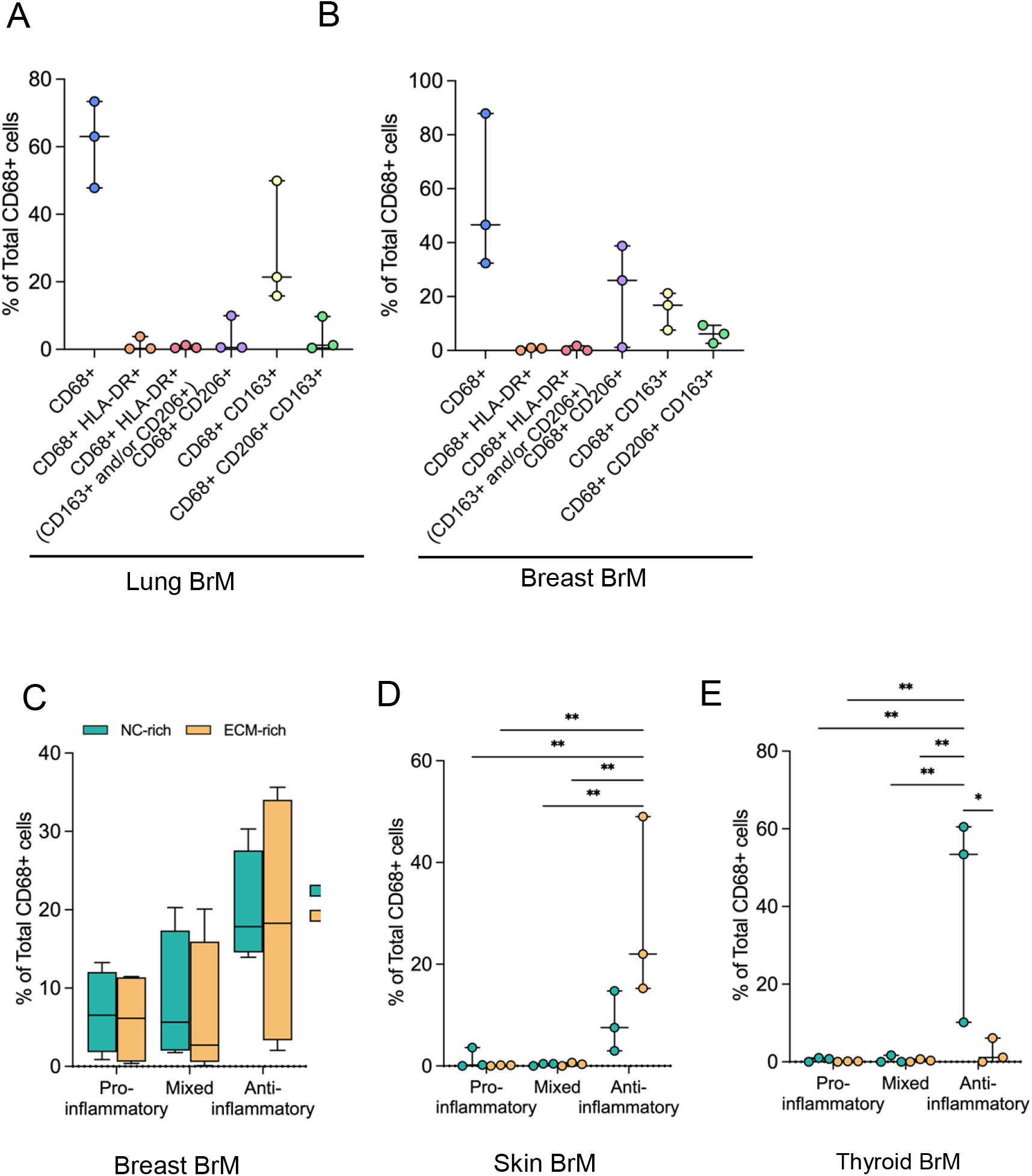
Localization of macrophage subsets in BrM. Percentage of macrophage subsets (CD68^+^, CD68^+^ HLA-DR^+^, CD68^+^ HLA-DR^+^ CD163^+^ and/or CD206^+^, CD68^+^ CD163^+^, and CD68^+^ CD206^+^ CD163^+^) as a proportion of total CD68^+^ cells in (**A**) skin (n=3) and (**B**) thyroid BrM (n=3) tissue. Statistical significance was determined by Kruskal-Wallis test, followed by Dunn’s test for multiple comparisons (α=0.05). Pro-inflammatory, anti-inflammatory and mixed macrophages as a proportion of total CD68^+^ cells (whole tissue core) in NC-rich and ECM-rich regions of (**C**) breast (n=5), (**D**) skin (n=3), and (**E**) thyroid BrM tissue (n=3). Statistical significance was determined by Wilcoxon matched-pairs signed rank test. Significance is represented by *(p<0.05), **(p<0.01), ***(p<0.001) and ****(p<0.0001).

**Figure S5.**
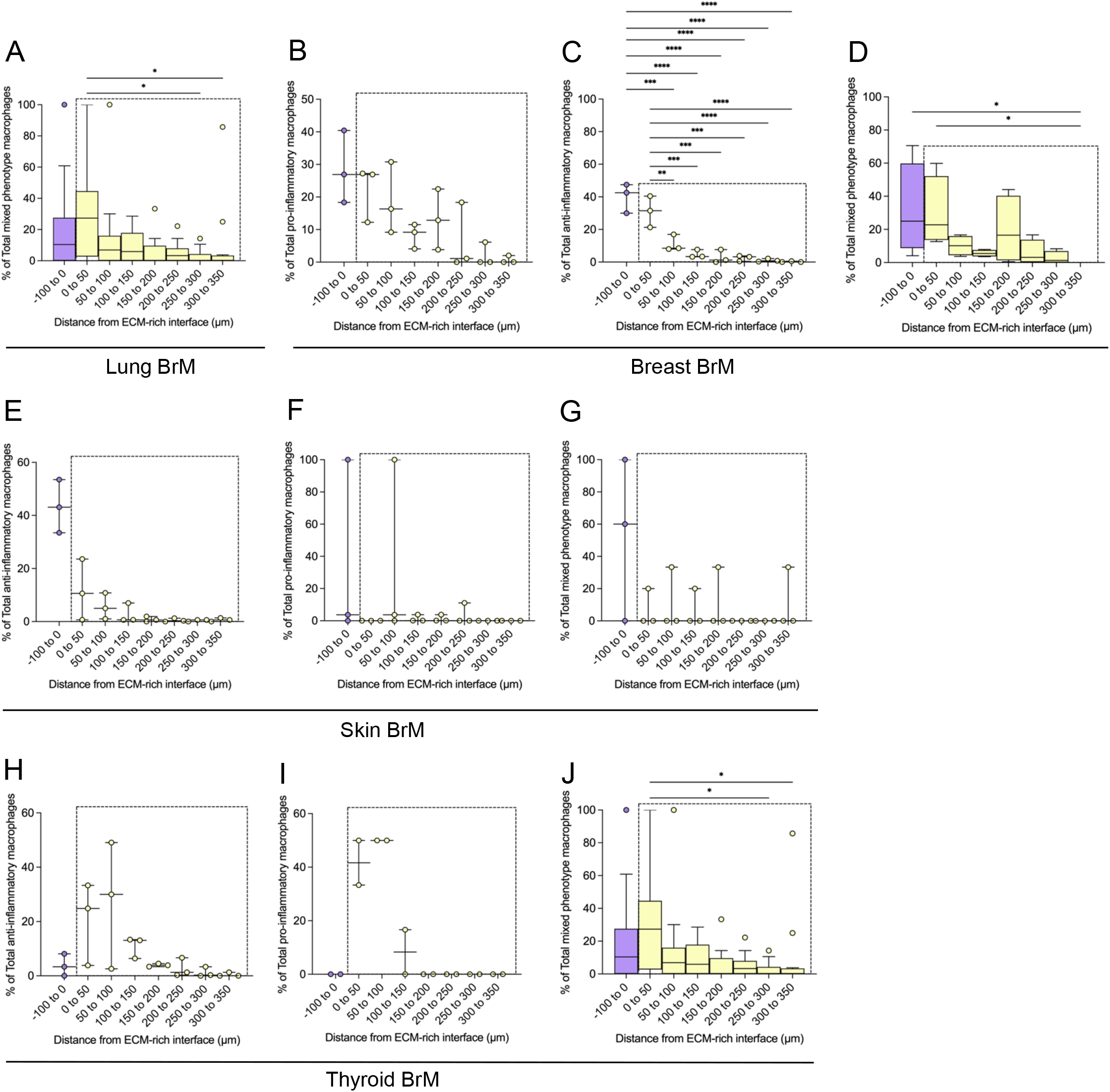
Infiltration analysis of macrophage subsets. Total anti-inflammatory (**C, E, H**), total pro-inflammatory (**B, F, I**), and total mixed (**A, D, G, J**) macrophage subset cell localization from the NC-rich and ECM-rich interface was determined (as a percentage of total CD68^+^ cells) and expressed as an infiltration distance from within the ECM-rich region (−100 to 0μm) into the NC-rich region (0 to 350μm). Data is shown for BrM from (**A**) lung (n=14), (**B-C**) breast (n=5), (**E-G**) skin (n=3), and (**H-J**) thyroid (n=3) primary tumor tissue. All results are presented as median +/− IQR. Statistical significance was determined by an ordinary one-way ANOVA. Significance is represented by *(p<0.05), **(p<0.01), ***(p<0.001) and ****(p<0.0001).

**Figure S6.**
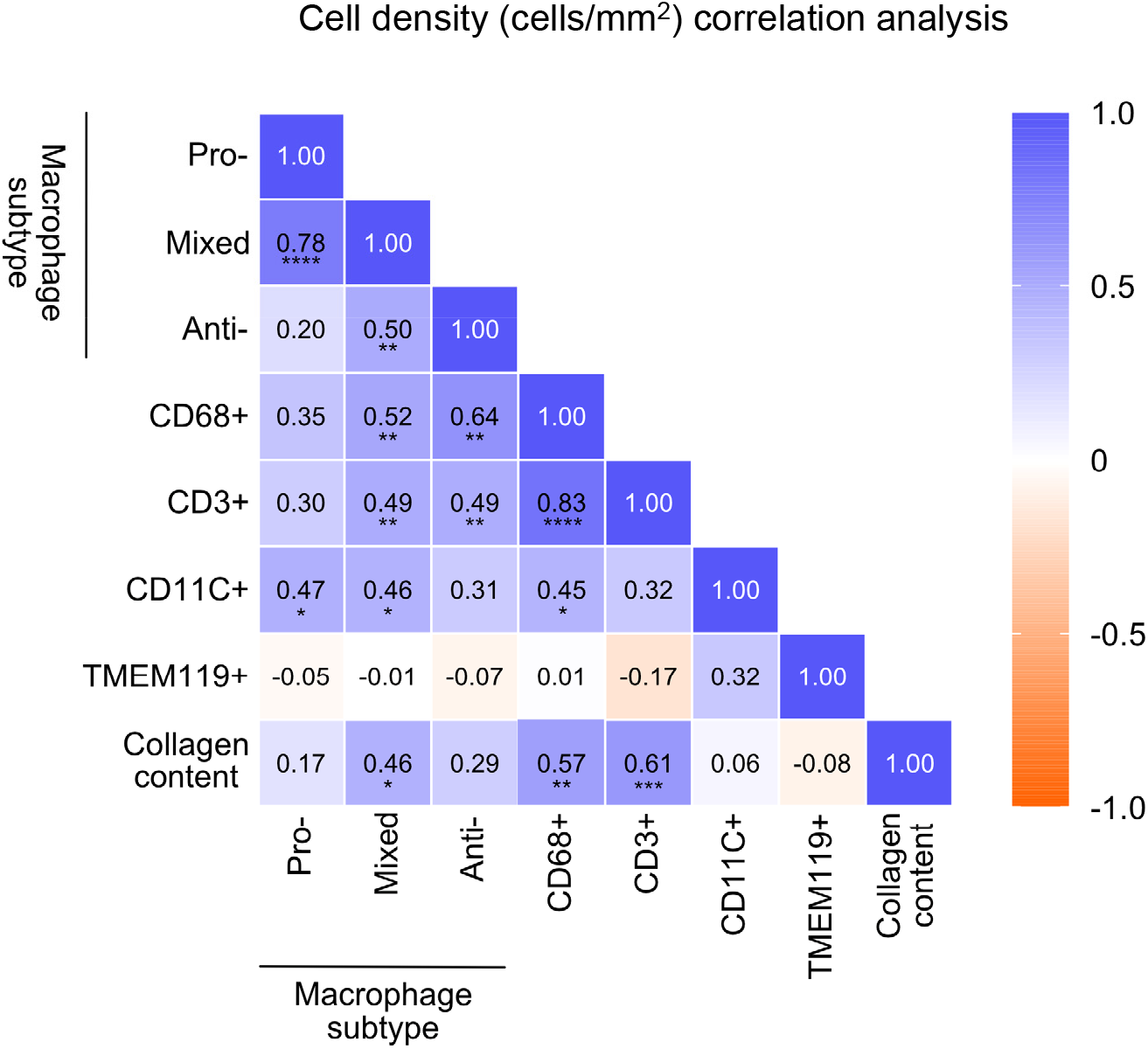
Collagen deposition correlates with the presence of macrophages and T-cells in BrM. Correlation heatmap between CD68^+^ (total CD68^+^), TMEM119^+^, CD11c^+^ and CD3^+^ cells, and pro-inflammatory, anti-inflammatory, and mixed macrophage subsets (cells/mm^2^) with collagen content in BrM tissue cores (n=28). Spearman rank correlation coefficients are indicated on the heatmap and significance is represented by *(p<0.05), **(p<0.01), ***(p<0.001) and ****(p<0.0001).

## References

1. Gavrilovic IT, Posner JB. Brain metastases: epidemiology and pathophysiology. J Neurooncol. 2005;75: 5–14.

2. Langley RR, Fidler IJ. The biology of brain metastasis. Clin Chem. 2013;59: 180–189.

3. McWilliams RR, Brown PD, Buckner JC, Link MJ, Markovic SN. Treatment of brain metastases from melanoma. Mayo Clin Proc. 2003;78: 1529–1536.

4. Galea I, Bechmann I, Perry VH. What is immune privilege (not)? Trends Immunol. 2007;28: 12–18.

5. Klemm F, Maas RR, Bowman RL, Kornete M, Soukup K, Nassiri S, et al. Interrogation of the Microenvironmental Landscape in Brain Tumors Reveals Disease-Specific Alterations of Immune Cells. Cell. 2020;181: 1643–1660.e17.

6. Friebel E, Kapolou K, Unger S, Núñez NG, Utz S, Rushing EJ, et al. Single-Cell Mapping of Human Brain Cancer Reveals Tumor-Specific Instruction of Tissue-Invading Leukocytes. Cell. 2020;181: 1626–1642.e20.

7. Seike T, Fujita K, Yamakawa Y, Kido MA, Takiguchi S, Teramoto N, et al. Interaction between lung cancer cells and astrocytes via specific inflammatory cytokines in the microenvironment of brain metastasis. Clin Exp Metastasis. 2011;28: 13–25.

8. Lin Q, Balasubramanian K, Fan D, Kim S-J, Guo L, Wang H, et al. Reactive astrocytes protect melanoma cells from chemotherapy by sequestering intracellular calcium through gap junction communication channels. Neoplasia. 2010;12: 748–754.

9. Novak U, Kaye AH. Extracellular matrix and the brain: components and function. J Clin Neurosci. 2000;7: 280–290.

10. Yoshida S, Takahashi H. Expression of extracellular matrix molecules in brain metastasis. J Surg Oncol. 2009;100: 65–68.

11. Virga J, Szemcsák CD, Reményi-Puskár J, Tóth J, Hortobágyi T, Csősz É, et al. Differences in Extracellular Matrix Composition and its Role in Invasion in Primary and Secondary Intracerebral Malignancies. Anticancer Res. 2017;37: 4119–4126.

12. Acerbi I, Cassereau L, Dean I, Shi Q, Au A, Park C, et al. Human breast cancer invasion and aggression correlates with ECM stiffening and immune cell infiltration. Integr Biol. 2015;7: 1120–1134.

13. Larsen AMH, Kuczek DE, Kalvisa A, Siersbæk MS, Thorseth M-L, Johansen AZ, et al. Collagen Density Modulates the Immunosuppressive Functions of Macrophages. J Immunol. 2020;205: 1461–1472.

14. Auvinen P, Tammi R, Parkkinen J, Tammi M, Agren U, Johansson R, et al. Hyaluronan in peritumoral stroma and malignant cells associates with breast cancer spreading and predicts survival. Am J Pathol. 2000;156: 529–536.

15. Liang Y, Xia W, Zhang T, Chen B, Wang H, Song X, et al. Upregulated Collagen COL10A1 Remodels the Extracellular Matrix and Promotes Malignant Progression in Lung Adenocarcinoma. Front Oncol. 2020;10: 573534.

16. Spivey KA, Banyard J, Solis LM, Wistuba II, Barletta JA, Gandhi L, et al. Collagen XXIII: a potential biomarker for the detection of primary and recurrent non-small cell lung cancer. Cancer Epidemiol Biomarkers Prev. 2010;19: 1362–1372.

17. Jansson M, Lindberg J, Rask G, Svensson J, Billing O, Nazemroaya A, et al. Prognostic Value of Stromal Type IV Collagen Expression in Small Invasive Breast Cancers. Front Mol Biosci. 2022;9: 904526.

18. Cohen JV, Kluger HM. Systemic Immunotherapy for the Treatment of Brain Metastases. Front Oncol. 2016;6: 49.

19. Priestley P, Baber J, Lolkema MP, Steeghs N, de Bruijn E, Shale C, et al. Pan-cancer whole-genome analyses of metastatic solid tumours. Nature. 2019;575: 210–216.

20. Bos PD, Zhang XH-F, Nadal C, Shu W, Gomis RR, Nguyen DX, et al. Genes that mediate breast cancer metastasis to the brain. Nature. 2009;459: 1005–1009.

21. Cosgrove N, Varešlija D, Keelan S, Elangovan A, Atkinson JM, Cocchiglia S, et al. Mapping molecular subtype specific alterations in breast cancer brain metastases identifies clinically relevant vulnerabilities. Nat Commun. 2022;13: 514.

22. Hebert JD, Myers SA, Naba A, Abbruzzese G, Lamar JM, Carr SA, et al. Proteomic Profiling of the ECM of Xenograft Breast Cancer Metastases in Different Organs Reveals Distinct Metastatic Niches. Cancer Res. 2020;80: 1475–1485.

23. Deasy SK, Erez N. A glitch in the matrix: organ-specific matrisomes in metastatic niches. Trends Cell Biol. 2022;32: 110–123.

24. Krakower C. Atlas of Human Glomerular Pathology: Correlative Light, Immunofluorescence, and UItrastructural Histology. JAMA. 1975;232: 1171–1171.

25. Papanicolaou M, Parker AL, Yam M, Filipe EC, Wu SZ, Chitty JL, et al. Temporal profiling of the breast tumour microenvironment reveals collagen XII as a driver of metastasis. Nat Commun. 2022;13: 4587.

26. Vennin C, Chin VT, Warren SC, Lucas MC, Herrmann D, Magenau A, et al. Transient tissue priming via ROCK inhibition uncouples pancreatic cancer progression, sensitivity to chemotherapy, and metastasis. Sci Transl Med. 2017;9. doi:10.1126/scitranslmed.aai8504

27. Junqueira LC, Bignolas G, Brentani RR. Picrosirius staining plus polarization microscopy, a specific method for collagen detection in tissue sections. Histochem J. 1979;11: 447–455.

28. Pekmezci M, Perry A. Neuropathology of brain metastases. Surg Neurol Int. 2013;4: S245–55.

29. Uhlen M, Zhang C, Lee S, Sjöstedt E, Fagerberg L, Bidkhori G, et al. A pathology atlas of the human cancer transcriptome. Science. 2017;357. doi:10.1126/science.aan2507

30. Uhlén M, Fagerberg L, Hallström BM, Lindskog C, Oksvold P, Mardinoglu A, et al. Proteomics. Tissue-based map of the human proteome. Science. 2015;347: 1260419.

31. Bowman RL, Klemm F, Akkari L, Pyonteck SM, Sevenich L, Quail DF, et al. Macrophage ontogeny underlies differences in tumor-specific education in brain malignancies. Cell Rep. 2016;17: 2445–2459.

